# Recognition of Necrosis and Ethylene-inducing like peptides confers disease resistance in *Brassica napus* and is modulated by *BSK1* in Arabidopsis

**DOI:** 10.1101/2023.08.01.551450

**Authors:** Hicret Asli Yalcin, Ricardo Humberto Ramirez-Gonzalez, Catherine N. Jacott, Burkhard Steuernagel, Gurpinder Singh Sidhu, Rachel Kirby, Emma Verbeek, Henk-jan Schoonbeek, Christopher J. Ridout, Rachel Wells

**Affiliations:** John Innes Centre, Norwich Research Park, Colney Lane Norwich, NR4 7UH, UK; TUBITAK Marmara Research Center, Natural Sciences, TUBITAK, 41470, Gebze, Kocaeli, Turkey; Genomics England, One Canada Square, E14 5AB, London, UK; Department of Microbiology, Faculty of Biology, University of Seville, Seville, Spain; University of East Anglia, Norwich Research Park, Norwich, NR4 7TJ, UK

## Abstract

Brassicas are important crops susceptible to significant losses caused by disease: thus, breeding resistant lines can mitigate the effects of pathogens. MAMPs (microbe-associated molecular patterns) are conserved molecules of pathogens that elicit host defence responses known as pattern-triggered immunity (PTI). Necrosis & Ethylene-inducing peptide 1-like proteins (NLPs) are MAMPs found in a wide range of phytopathogens, including major disease-causing fungal species. We studied the response to the BcNEP2 from *Botrytis cinerea* as a representative NLP in *Brassica napus* to improve our understanding of recognition mechanisms that could enable the development of disease-resistant crops.

To genetically map regions responsible for NLP recognition, we used an associative transcriptomics (AT) approach using diverse *B. napus* accessions and bulk segregant analysis (BSA) on DNA pools created from a bi-parental cross of NLP-responsive (Ningyou1) and non-responsive (Ningyou7) lines*. In silico* mapping with AT identified two peaks associated with NLP recognition on chromosomes A04 and C05 whereas the BSA narrowed it down to a main peak on A04. BSA delimited the region associated with NLP-responsiveness to 3 Mbp, containing ∼245 genes on the Darmor*-bzh* reference genome. Variants detected in the region were used for KASP marker design and four KASP markers were identified co-segregating with the phenotype. The same pipeline was performed with the ZS11 genome, and the highest associated region was confirmed on chromosome A04. Comparative BLAST analysis revealed there were unannotated clusters of RLP homologs on ZS11 chromosome A04. To reduce the number of candidate genes responsible for NLP recognition, RNA-Seq data was used to detect the unannotated expressed putative genes. Screening the BSA Ning1×7 population demonstrated a highly significant association between NLP-recognition and resistance to *Botrytis cinerea*. Also, the lines non-responsive to NLP had significantly greater response to the bacterial MAMP flg22. Additionally, *BnaA01g02190D*, a homologue of Arabidopsis *AtBSK1* (*At4g35230*) *BR-SIGNALLING KINASE1,* was associated with a high BcNEP2-induced ROS response phenotype. We show that in Arabidopsis, *Atbsk1* mutants had significantly lower response to BcNEP2 and increased susceptibility to *B. cinerea* (*p-value*=1.12e-14***). Overall, the results define the genomic location for NLP-recognition on the *B. napus* genome and demonstrate that NLP recognition has a positive contribution to disease resistance which can have practical application in crop improvement.

## Introduction

Necrosis- and ethylene-inducing like proteins (NLPs) are a superfamily of proteins widely found in bacteria, fungi, and oomycetes (Seidl and Van den Ackerveken, 2019). NLPs can be present in microorganisms that have a plant-pathogenic lifestyle, but they also occur in non-pathogenic taxa suggesting their role is not limited to virulence in plants (Bhatti et al., 2017; Tekaia and Latgé, 2005). NLPs can contain a microbe-associated molecular pattern (MAMP) motif that activates pattern-triggered immunity (PTI) in plants leading to defence gene induction, production of reactive oxygen species (ROS) and other protective mechanisms. Recognition of the NLP necrosis and ethylene-inducing like peptide 2 (BcNEP2) by RECEPTOR-LIKE PROTEIN 23 (AtRLP23) contributes to some resistance against *Botrytis cinerea* in *Arabidopsis thaliana* (Ono et al., 2020). In translational research studies, silencing of *BcNEP2* in *B. cinerea* and its corresponding homologue *SsNEP2* in *Sclerotinia sclerotiorum* by dsRNA resulted in a significant reduction of lesion size in *B. napus* (McLoughlin et al., 2018). However, BcNEP2-recognition could not be associated with resistance to *B. cinerea* in 12 *B. napus* accessions possibly because of their diverse genetic backgrounds and the limited extent of the study (Schoonbeek et al., 2022).

In addition to NLPs, other MAMPs including chitin, flagellin and elongation factor Tu (EF-Tu) are well-studied in the Brassicaceae family. Short amino acid peptides of EF-Tu (elf18) can induce PTI in the Brassicaceae whereas flagellin (flg22) and chitin can induce PTI in many other plant species. We previously described methods to evaluate PTI in *B. napus* (Lloyd et al., 2017, Lloyd et al., 2014). MAMPs are recognised by pattern recognition receptors (PRRs), which include receptor like kinases (RLKs) and receptor like proteins (RLPs) and may interact with co-receptors to activate defence responses. RLP23 recognises NLPs and forms a constitutive complex with the LRR receptor kinase, SOBIR1, in a ligand-independent way and also recruits a second LRR receptor kinase, BAK1 into a complex upon ligand binding (Albert et al., 2015). The leucine-rich repeat (LRR)-RLK protein family XII includes the PRRs FLAGELLIN SENSITIVE 2 (FLS2) and ELONGATION FACTOR (EF)-Tu RECEPTOR (EFR) in Arabidopsis which respectively recognise flg22 and elf18 (Felix et al., 1999; Gomez-Gomez and Boller, 2000, Kunze et al., 2004; Zipfel et al., 2006). The FLS2 receptor has previously been shown to physically associate with a co-receptor BSK1 in Arabidopsis during the activation of immunity (Shi et al., 2013). However, the role of BSK1 in NLP recognition is not known.

The Brassicaceae is a large plant family of 338 genera and more than 3709 species (Schmidt et al., 2001). Six species of major agricultural and horticultural importance in the genus *Brassica* are defined by their genome arrangement (A, B or C) for diploid species, *B. rapa* (AA), *B. nigra* (BB) and *B. oleracea* (CC), that are hybridised in the allotetraploid (amphidiploid) species, *B. juncea* (AABB), *B. napus* (AACC) and *B. carinata* (BBCC) (commonly referred to as polyploids). The subject of this study, *B. napus*, is the most important temperate oilseed crop in the world with a production of close to 70 million tonnes (Raboanatahiry et al., 2021). *B. napus* is affected by a range of pathogens which affect yield and quality of the crop and improved methods are needed for disease control. We previously used *B. cinerea* as a model pathogen with extensive resources to translate disease resistance research from Arabidopsis to *B. napus*. We also showed that some *B. napus* accessions could recognise NLPs from various crop pathogens such as *B. cinerea* (BcNEP1 and BcNEP2), *Verticillium longisporum* (VlNLP1), *Leptosphaeria maculans* (LmNLP1), *Alternaria brassicicola* (AbNLP2) and also *Hyaloperonospora arabidopsidis* (HaNLP3) which, if this can contribute to disease resistance, could provide a new approach to disease control (Schoonbeek et al., 2022).

The polyploid nature of the *B. napus* genome results in multiple homologues of candidate receptor and co-receptor genes involved in NLP recognition when compared to Arabidopsis (Schoonbeek et al., 2022). Although this complexity presents challenges for polyploid species, the bioinformatics resources for gene identification are well advanced for *B. napus* including the availability of two reference genomes, ZS11 (Song et al., 2020) and Darmor-*bzh* (Chalhoub et al., 2014). Genome-wide association studies (GWAS) and associative transcriptomics (AT) involve associating single nucleotide polymorphisms (SNPs) in genomes or transcriptomes respectively with polymorphic traits in genetically diverse accessions and have been used to identify candidate resistance gene loci for *S. sclerotiorum* (Sclerotinia stem rot), *Plasmodiophora brassicae* (clubroot) and *Pyrenopeziza brassicae* (light leaf spot) in *B. napus* (Dakouri et al., 2021; Fell, Muthayil Ali, et al., 2022; Roy et al., 2021).

Bulked segregant analysis (BSA) is a technique used for identifying the genetic markers for specific target loci responsible for interesting traits (Michelmore et al., 1991). The advantages of BSA are that candidate genes can be identified quickly because heterozygous plants are sufficient to map the corresponding region and, since bulked samples are used for sequencing, the costs are reduced (Zou et al., 2016). The method is also very flexible since any segregating population can be used and any NGS-enabled protocol can be applied such as RNA-Seq (Trick et al., 2012) or whole-genome re-sequencing (Tudor et al., 2020). Already, many agronomically important traits have been investigated using BSA with different crop species including *Oryza sativa, Triticum aestivum L.* and *B. napus* (Ramirez-Gonzalez, Segovia, et al., 2015; Takagi et al., 2013; Wang et al., 2016). One of the most recent BSA studies in *B. napus* successfully identified a major QTL at a 10Mbp region on chromosome A02, which includes orthologues of *AtFLC*; *FLOWERING LOCUS C* (*BnaFLC.A02*) and *AtFT*; *FLOWERING LOCUS T* (*BnaFT.A02*) (Tudor et al., 2020).

Our study is based on the hypothesis that NLP recognition contributes to disease resistance in *B. napus* in a similar manner to that in Arabidopsis. To address this, we created a mapping population from NLP-responsive and non-responsive lines and used this to test resistance against *B. cinerea*. To localise the candidate NLP receptor and potentially co-receptors, we used a combination of AT with a *B. napus* diversity set and a more targeted BSA approach from pools of NLP-responding and non-responding progeny lines of the mapping population. Our strategy and results have potential applications in improving disease resistance in *B. napus*.

## Material & Methods

### Plant material

A *B. napus* diversity panel of 189 lines from the Renewable Industrial Products from Rapeseed (RIPR) panel was used for AT analysis (Havlickova et al., 2018). Genotypes were split into four batches for sampling and two experimental replicates were performed.

For BSA analysis, reciprocal crosses between semi-winter oilseed rape (OSR) accessions Ningyou1 and Ningyou7 (Source, OCRI, Wuhan, China) were generated by manual crossing with two replicates. Tapidor, a winter OSR accession, from the *B. napus* diversity set and Brassica Germplasm Collection at the John Innes Centre, UK, was used as a control accession. Parental lines for crossing and F_1_ plants for seed production were sown in Levington F2 with grit and grown in a lit glasshouse with a 16-hour photoperiod at 18/12°C Day/night temperatures (∼6 months). For disease and MAMP assays, plants were grown in Levington F2 with grit for 4-5 weeks in controlled environment rooms (CERs) at 22°C, 70% relative humidity, under 10/14 hours day/night cycle.

Arabidopsis plants were sown in Levington F2 with 15% 4mm grit in 24-cell trays. Seeds were stratified in the dark for two days at 4-5°C and trays were covered with a transparent plastic lid to increase the humidity for the first 1-2 weeks. Following stratification, plants were grown at 20-22/18-20°C Day/night, 70% relative humidity, under 10/14 hours day/night cycle in a CER for 5-6 weeks prior to phenotyping. Arabidopsis *Atbsk1-1* mutant seeds and Col-0, used as wild type, were kindly obtained from Prof. Dingzhong Tang, Chinese Academy of Sciences, Institute of Genetics and Developmental Biology (Nie et al., 2011).

### Reactive Oxygen Species (ROS) Assay

#### Microbe associated molecular pattern (MAMP) peptides

Peptides of flg22 (22 amino acid long flagellin fragment; QRLSTGSRINSAKDDAAGLQIA, Peptron http://www.peptron.co.kr, Korea, dissolved in sterile H_2_O at 10mM) and BcNEP2 (*Botrytis cinerea* Necrosis and Ethylene-inducing protein 2; AIMYSWYMPKDEPSTGIGHRHDWE, Genscript www.genscript.com, at 87.5% purity and dissolved in DMSO at 10mM) were used for detection of oxidative burst. Peptides were aliquoted to 100µM in H2O and stored at −20°C before use.

#### Detection of oxidative burst

A luminol/peroxidase based assay was used to measure MAMP-induced reactive oxygen species (ROS) production (Lloyd, 2014). 4mm leaf discs from *B. napus* (3^rd^ leaf) or Arabidopsis were incubated in 200µl of sterile water in a 96-well plate overnight, in the dark, at room temperature. The water was drained and replaced by 100µl solution containing; 0.2nM luminol at 34mg/l, horseradish peroxidase (HRP) at 20 mg/l (Sigma), and the selected MAMP at concentrations of 50nM, 10nM or 2nM. Luminescence was recorded over a 40-minute period and displayed as photon production quantitated as relative light units (RLUs) over this period. Emitted photons were counted using a Varioskan Flash plate reader (ThermoFisher Scientific, Waltham, MA, USA).

### Botrytis cinerea disease assays

*Botrytis cinerea* strain B05.10 (Schoonbeek et al., 2001) was grown on 1/5 Potato Dextrose Agar (PDA) or on MEYAA plates including; Malt Extract Agar (MEYA, OXOID# CM0059) at 30g/l with Yeast extract at 2g/l and Agar (FORMEDIUM) at 5g/l at 21°C with no selection. *B. cinerea* plates were grown for at least 2 weeks before collecting spores for inoculation.

One day prior to the inoculation of *B. napus* leaf discs, 200µl from *B. cinerea* spores with 2.5×10^6^ spores/ml concentration were spread on 1/5 PDA plates and grown at 21°C. Leaf discs (22mm, single disk/plant, up to 16 replicates) cut from the 4^th^ leaf of *B. napus* plants were placed on 0.6% water agar plates in square Petri dishes and left in the dark, overnight. The next day, leaf discs were inoculated with *B. cinerea* plugs (4mm diameter) and incubated in the growth cabinet at 21°C with a relative humidity of 85-100% and at low light condititons with photon flux density in between 10-20µmol m^-2^ s^-1^. The lesion size was measured after 3 days with a digital caliper.

Prior to inoculation of Arabidopsis plants, *B. cinerea* spores were diluted to 2.5×10^5^ spores/ml concentration in 1⁄2 Potato Dextrose Broth (PDB) solution and shaken at room temperature for 90 minutes. 5μl droplets from the spore solution were placed on selected 5-6 leaves of individual plants grown in 24-cell trays. Trays were covered with plastic lids sealed with parafilm and transferred to a growth cabinet at 21°C with a relative humidity of 85-100% and low light (10-20µmol m^-2^ s^-1^). Lesion sizes were measured after 3 days with a digital caliper.

### Associative Transcriptomics (AT)

Total RLU over 40 minutes for each genotype was calculated using a linear fixed- and random effects model with *B. napus* genotype as a fixed effect and experimental replicate and batch as random effects. Estimated marginal means were calculated for each *B. napus* genotype using *emmeans* (version 1.8.0) (R package). These data were transformed into binary values (1 or 0, for responding and non-responding lines, respectively) for AT analysis.

The RIPR genotype (SNP) and expression level datasets (Havlickova et al., 2018) were obtained from York Knowledgebase (http://yorknowledgebase.info). This dataset was reduced to only include accessions used in this study. Our recently updated population structure (Fell, Ali, et al., 2022) was used for the AT analysis.

Mapping was performed using an R based pipeline (Nichols, 2022) using GAPIT Version 3 (Lipka et al., 2012; Wang and Zhang, 2021). Analyses were conducted using a generalised linear model (GLM); Bayesian-information and Linkage-disequilibrium Iteratively Nested Keyway (BLINK) (Huang et al., 2019), and Fixed and random model Circulating Probability Unification (FarmCPU) (Liu et al., 2016) to determine the optimal model. The false discovery rate (FDR) was determined using the Shiny implementation of the q-value R package (Storey, 2011).

Linkage disequilibrium (LD) varies between and across chromosomes. To determine specific marker LD we calculated the mean pairwise R^2^ for the peak marker to all markers on the chromosome using TASSEL Version 5.0 using the site by all analysis option (Bradbury et al., 2007). Markers were considered in LD when R^2^> 0.2.

### Bulk Segregant Analysis (BSA)

#### Creation of F_2_ segregant population

Five seeds from each Ningyou1 and Ningyou7 F_1_ cross were sown and leaf samples from each F_1_ plant were used for DNA isolation with the DNeasy® Plant kit (Qiagen), according to the manufacturer’s protocol. Publicly available microsatellite markers (http://www.brassica.info/resource/markers/ssr-exchange.php) were used to detect polymorphic markers between the parental accessions. Polymorphic marker, sR12095, **(Table S1, Figure S1)** was used to confirm F_1_ crosses with a heterozygous genotype. PCR with SSR sR12095 (Agriculture and Agri-Food Canada AAFC) was conducted by using AmpliTaq Gold™ DNA Polymerase (ThermoFisher) in a final reaction volume of 20μl with 2µl of the diluted genomic DNA (100ng), 0.5μM from each primer pairs, 2µl PCR buffer, 1 units AmpliTaq Gold™ DNA Polymerase (5u/µl), 2.5mM dNTPs and water to 20μl. The optimized PCR programme had an initial denaturation step at 94°C for 10 minutes with a PCR protocol followed as 8 cycles of 94°C for 15 seconds, 50°C for 15 seconds and 72°C for 30 seconds. This was followed with a PCR protocol beginning with 32 cycles of 94°C for 15 seconds, 50°C for 15 seconds, 72°C for 30 seconds and ending with a final cool down at 8°C. DNA bands visualized on 3.0% Agarose gel by using AlphaImager EP system. Each plant was also phenotyped for ROS production by 50nM BcNEP2 and 20nM flg22. Successfully crossed F_1_ plants were selfed to develop the Ningyou1 x Ningyou7 (Ning1×7) F_2_ population. 960 F_2_ seeds were sown and grown under controlled environment room (CER) conditions for ROS phenotyping for BSA.

#### Experimental design - creation of bulked DNA pools

BcNEP2-induced ROS production data from the F_2_ population was normally distributed (Figure 2). Extremes from the tails of the normal distribution representing ∼5% of the population (30 of 747 NLP-responder lines) were selected to create DNA pools with the phenotypic features as shown in Figure S3 and Table 1.

**Table 1:** The phenotypical features of the selected individuals from F_2_ population for the BSA pools.

| Phenotype | Pool1 | Pool2 | Pool3 | Pool4 |
| --- | --- | --- | --- | --- |
| Magnitude of flg22-induced ROS | Low | High | Low | High |
| BcNEP2 recognition (-/+) | - | - | + | + |
| Magnitude of BcNEP2-induced ROS | - | - | Low | High |

**Note:** Individual F<sub>2</sub> plants were selected based on their magnitude of BcNEP2- and flg22-induced ROS production and their ability to recognize BcNEP2 molecule. “Low” MAMP-induced ROS production defines individuals with ROS production lower than 1.0E5 RLU. “High” MAMP-induced ROS production defines individuals with ROS production higher than 8.0E5 RLU. - Classification of the individual plants according to their ability to recognize BcNEP2 molecule is defined as “-” for blind individual plants and “+” for the responders.

To isolate high-quality DNA from the *B. napus* leaves for sequencing, leaf tissue from individual F_2_ plants was collected and frozen at −80°C using liquid Nitrogen. 0.05g leaf samples from each selected line were bulked for grinding to create pools. To ensure the same quantity of samples used for each pool and the parental samples, 1.5g of leaf material was used for parental lines. DNA was isolated using an improved CTAB-based method with extra Chloroform/Isoamyl Alcohol and Phenol/Chloroform/Isoamyl Alcohol steps and use of NaAc solution for precipitation (Tudor et al., 2020).

### Sequencing & Bulk Segregant Analysis

Illumina sequencing (350bp insert DNA library, 150-bp paired-end reads, 40x coverage, via Illumina NovaSeq PE150 Platform) was performed for the bulked pools and parents. Sequence quality was evaluated with FastQC v0.11.8 (https://www.bioinformatics.babraham.ac.uk/projects/fastqc/). Reads were aligned to *B. napus* reference genomes Darmor-*bzh* (Chalhoub et al., 2014) and Zhongshuang11 (Song et al., 2020) using *bwa* v0.7.1 (Li and Durbin, 2010). The resulting SAM files were converted to BAM files using *samtools* v1.9 (Danecek et al., 2021), and all BAM files containing data from the same pool were merged. After indexing, SNP calling was performed per chromosome using *freebayes* v1.1.0.46 (Garrison and Marth, 2012). The Variant Call Format (VCF) files were concatenated using *bcftools* v1.8 (Danecek et al., 2011). Filtering was applied using *bcftools-1.8*. to exclude variations with quality lower than 2000, depth lower than 20 and those coming from Darmor-*bzh.* Markers were named according to the reference genome (“BnDar” for Darmor-*bzh* and “BnZS” for Zhongshuang11), chromosome/scaffold type (A or C) and number (01 to 10), genomic position on the related chromosome and sequence of the alternative variant.

The ratio between the frequency of each informative base at each variant position for the pools was calculated. For variations with an informative base derived from a responsive background, the Bulk Frequency Ratio (BFR) was calculated by dividing the frequency in the responsive bulk by the frequency in the non-responsive bulk (Trick et al., 2012). The formula for calculation of the BFR values for each variant is (Counts of Alternative (Alt) variation is called as AO, counts of Reference (Ref) variation is called RO and count of total reads (Depth) is called DP in the formula):

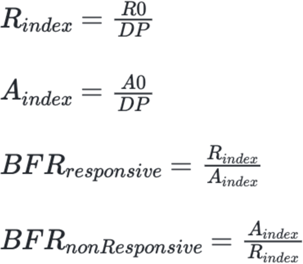

The *ggplot2* package (Wickham, 2016) was used to create the Manhattan plots with BFR values of each variant anchored to the chromosomes.

*SnpEff*, a functional effect prediction toolbox, was used to predict the effects of genetic variants on genes and proteins (Cingolani, Platts, et al., 2012). Related information from VCF files were obtained through *SnpSift* (Cingolani, Patel, et al., 2012). To prepare the input format for futher filtering, VCF files were converted to TXT file format by using *bcftools* v1.8. Then, BioMart database (Durinck et al., 2005, 2009) was used to identify any possible Arabidopsis homologues and coded domain features of the *B. napus* genes.

All the code used in BSA is available on the github repository: https://github.com/hicretyalcin/BSA_Brassica_napus.

### KASP Assay

Kompetitive allele specific PCR (KASP) markers were developed by using PolyMarker (Ramirez-Gonzalez, Uauy, et al., 2015). A 200bp genomic sequence, including the SNP, was generated to start the pipeline. Designed primers included standard FAM or HEX compatible tails (FAM tail: 5’ GAAGGTGACCAAGTTCAT-GCT 3’; HEX tail: 5’ GAAGGTCGGAGTCAACGGATT 3’ (Sigma)) **Table S2)**. DNA was isolated from samples with known phenotypes from the F_2_ population with DNeasy® Plant kit (Qiagen) according to the manufacturer’s protocol. The DNA plate contained 21 lines from: each bulked Pool (Pool1, Pool2, Pool3, and Pool4); 3 individual plant DNA samples from Ningyou1; 3 individual plant DNA samples from Ningyou7; and DNA samples of 6 different NLP-responsive *B. napus* lines (Swu8, Yudal, Chuanyou2, Tribune, Taisetsu, and ZS11) respectively.

For the KASP Assay, DNA was diluted to 10ng/μl, then 1.8μl of DNA was dispensed into 384 well PCR plate. To obtain a DNA pellet, the PCR plate then dried in an incubator. Then, PCR mastermix containing 1.2μl water, 1.2μl PACE® (3CR Bioscience, UK) mix and 0.03μl primer mix per well was dispenced to the wells. Primer mix was made up with 46μl water, 12μl of each forward primer, and 30μl of the common reverse primer. The PCR programme was 94°C for 15 minutes followed by 10 touchdown cycles at 94°C for 20 seconds and 65°C for 1 minute, dropping by 0.8°C per cycle to 57°C. Then another 45 cycles at 94°C for 20 seconds and at 57°C for 1 minute. The plate was then read on a BMG PHERAstar plate reader and scored using LGC’s *Klustercaller* program.

### Genetic Map construction

Linkage analysis was made by using R version 3.6.1 with the *qtl* package version 1.46-2 (Broman, 2010). Genetic distances of each marker to the phenotype based on recombination frequency was calculated.

### Detection of Expressed Putative Genes on ZS11 Genome

To find novel gene candidates that have previously been missed by gene annotation of ZS11, RNA-Seq data from ZS11 was aligned to the genome sequence (Song et al., 2020) using hisat2 (Kim et al., 2019), version 2.1.0 with standard parameters. Resulting SAM files were sorted and converted to BAM format using samtools sort (Danecek et al., 2021), version 1.10. Resulting BAM files were indexed using samtools index. Regions of interest in the genome were manually inspected using Integrative Genomics Viewer (IGV) (Robinson et al., 2011). *A. thaliana* homologues were assigned by reciprocal BLAST search strategy. Gene IDs obtained as hits after performing BLAST search (e-value cut off 1e-5) against Darmor-*bzh* v10 using TAIR genome as query, were used as query genes to run a reciprocal blast search with TAIR genome as target (e-value cut off 1e-5). Genes were marked as mapped if the reciprocal blast query Darmor-*bzh* gene returned the same *A. thaliana* gene as the highest scoring hit with the as the original Darmor-*bzh* gene with an e-value < 1e-5 in the previous search.

### Genotyping of *A. thaliana bsk1-1* mutants

DNA from Arabidopsis leaf samples was extracted with DNeasy® Plant kit (Qiagen) according to the manufacturer’s protocol. All samples were verified to contain the *bsk1-1* mutation using the polymerase chain reaction (PCR) with cleaved amplified polymorphic sequence (CAPS) primers (Shi et al., 2013) **(Table S1)** followed by a digestion with BsuRI (HaeIII) enzyme. DNA bands visualized on 2.4% agarose gels.

### Quantification of gene expression in *A. thaliana*

#### MAMP pre-treatment

For MAMP treatment, 100 nM BcNEP2 was dissolved in water with 0.01% DMSO prepared and injected to at least 6 leaves of individual 5-week old Col-0 and *bsk1-1* plants from the treatment group. Plants belonging to the control (Mock) group were injected with Mock solution (Water with 0.01% DMSO). Each leaf was injected with ∼200µl solution using a 1ml syringe. Samples were collected at 0h, 12h, and 24h and frozen in −80°C using liquid nitrogen. Additionally, for no-Treatment, samples were collected from 3 Col-0 and 3 *bsk1-1* plants. At least 3 biological replicates were performed for each treatment. 3 technical replicates were used from each biological replicate.

#### Quantitative Reverse Transcription Polymerase Chain Reaction (qRT-PCR)

Frozen samples were ground into a fine powder using liquid Nitrogen. RNA was extracted with the RNeasy^®^ plant mini kit (QIAGEN) and, after isolation, samples were treated with DNAse (TURBO DNA-*free*^TM^ kit (INVITROGEN)), according to the manufacturer’s protocols. To synthesize the cDNA, SuperScript IV^TM^ Reverse Transcriptase (INVITROGEN) was used according to the manufacturer’s protocol with 1μg of DNase-treated RNA/20μl reaction.

Primers were designed using Primer3Plus software (Untergasser et al., 2007). At least two primer pairs were designed for each gene of interest and obtained from Sigma. Only primer pairs with amplification efficiencies over 90-100% were selected for qRT-PCR analysis **(Table S1)**. Each of the reactions was performed with 12μl reaction mix containing 1:20 diluted cDNA. SYBR® Green JumpStart^TM^ Taq from Sigma-Aldrich was used according to the manufacturer’s protocol. The qRT-PCR programme was 96°C for 4 minutes, followed by 45 cycles at 94°C for 19 seconds, 60°C for 19 seconds and 72°C for 22 seconds. Cycle threshold (CT) values were used to determine the relative transcript levels according to the 2ΔCT method using *ACTIN* as a reference (Shi et al., 2013). To calculate the fold induction, transcript levels of the genes in BcNEP2-treated samples were compared with Mock treated controls.

## Results

### 1. Associative Transcriptomics identifies a region associated with NLP-recognition on chromosome A04

To determine the phenotypic variation in NLP recognition in *B. napus* genotypes, we measured ROS production after treatment with BcNEP2. The majority of *B. napus* lines did not respond to BcNEP2, with only 12 out of 189 lines responding **(Table S3)**. These data suggested that BcNEP2 recognition is a qualitative trait and thus, can be represented as binary values (1 or 0, for responding and non-responding genotypes, respectively) **(Figure 1b, Table S4)**.

**Figure 1.**
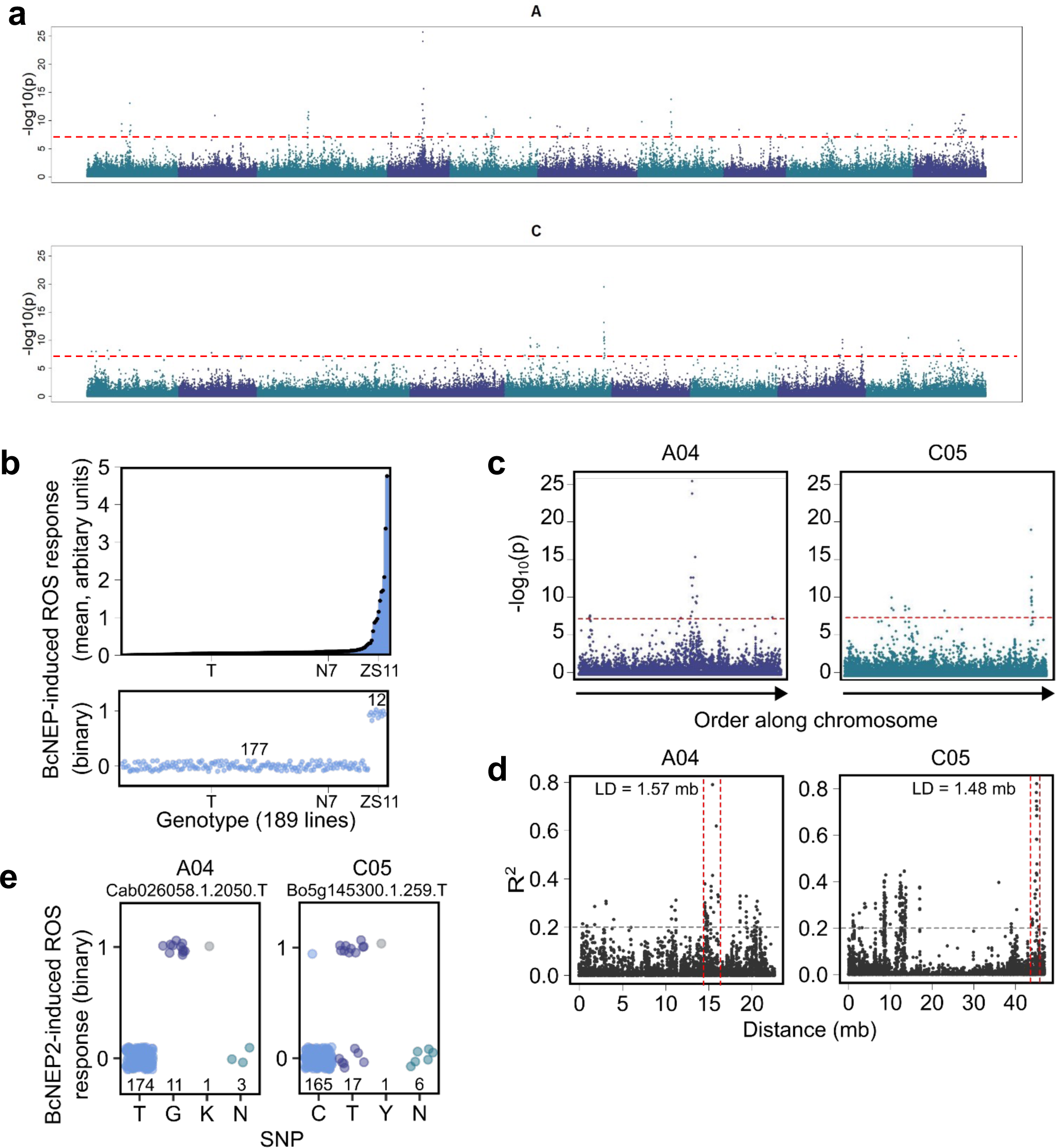
Associative Transcriptomic analysis of BcNEP2-induced ROS production in 191 *B. napus* genotypes. **(a)** Manhattan plots showing marker-trait association for binary ROS production to BcNEP2. Red line indicates FDR <0.05. **(b)** Mean (calculated using *eemeans* package in R) and binary ROS production of *B. napus* genotypes to BcNEP2. Reference genotypes Tapidor (T), Ningyou 7 (N7) and Zhongshuang11 (ZS11) are indicated. **(c)** Manhattan plots showing marker-trait association for binary ROS production. The X-axis indicates SNP location by order along chromosome; the y-axis indicates the −log10(p) (p-value). Red lines indicate FDR <0.05. **(d)** Linkage disequilibrium of the highest associating marker from peaks on chromosome A04 (Cab026058.1.2050.T) and C05 (Bo5g145300.1.256.T). Grey line indicates R^2^ value of 0.2, red lines indicate the area of LD. **(e)** Segregation of BcNEP2-induced ROS production in the *B. napus* genotypes of Cab026058.1.2050.T (A04) and Bo5g145300.1.256.T (C05).

To identify loci associated with NLP response, we performed AT analysis using binary values. AT analysis indicated two major peaks, on chromosomes A04 and C05 **(Figure 1a, 1c)**. Assessment of phenotypic variation segregating with alleles for the highest marker at the A04 peak, Cab026058.12050.T, revealed that all accessions inheriting a “G” at this locus (11 accessions) could respond to BcNEP2, and all accessions inheriting a “T” (174 accessions) could not **(Figure 1e. Table S4)**. On the otherhand, for the highest marker at the C05 peak, Bo5g145300.1.259.T, only 10 out of 17 accessions inheriting a “T” at this locus could respond to BcNEP2, and 164 out of 165 accessions inheriting a “C” could not **(Figure 1e)**.

We calculated the genetic distance in linkage disequilibrium (LD) with Cab026058.12050.T (A04) and found a region of 1.57 mb **(Figure 1d)** containing 250 genes in the *B. napus* pantranscriptome (**Table S5**). We calculated 1.48 mb in LD with Bo5g145300.1.259.T (C05) **(Figure 1d)**, which corresponded to 252 genes in the *B. napus* pantranscriptome **(Table S6)**. We performed BLASTP analysis to determine the putative *A. thaliana* orthologs of the *B. napus* genes in these two regions and found that these associations were not in homeologous genomic regions.

### 2. A single dominant gene locus is associated with NLP-recognition within the Ningyou1 x Ningyou7 population

To define the responsible gene or associated regions on *B. napus* genome, we developed a mapping population was developed between Ningyou1 (NLP-responsive) and Ningyou7 (NLP-non-responsive), henceforth known as Ning1×7. All F_1_ plants from the Ning1×7 cross were heterozygous and responsive to BcNEP2 **(Figure S1, S2)**.

F_1_ plants were selfed and F_2_ seeds obtained from them. The ROS production by 50nM BcNEP2 in responding F_2_ individual plants ranged from ∼ 2.6×10^3^ to 1.3×10^6^ RLU **(Figure 2a)**. 178 out of 925 F_2_ individuals produced no ROS response following BcNEP2 treatment, representing ∼19% of the population. This ratio was expected to be ∼25% (3:1, responder: non-responder) if a single dominant gene model (*X^2^*-test; *P*<0.001) is predicted. Although the ratio obtained is close to that model, the slight difference between predicted and observed values could result from seedling selection during the transplanting process. Specifically, during the phenotyping process, it was observed that some of the non-responding individuals were smaller and had chlorosis and could have been selected against for vigour during transplantation. Since all heterozygote F_1_ individuals derived from at least one NLP-responsive parent, our combined results indicate that a single dominant gene locus is responsible for NLP-recognition. All individuals from the cross could mount a ROS response to 20nM flg22 to some extent, confirming that the cellular mechanisms for PTI were functional in NLP non-responders **(Figure 2c)**.

**Figure 2.**
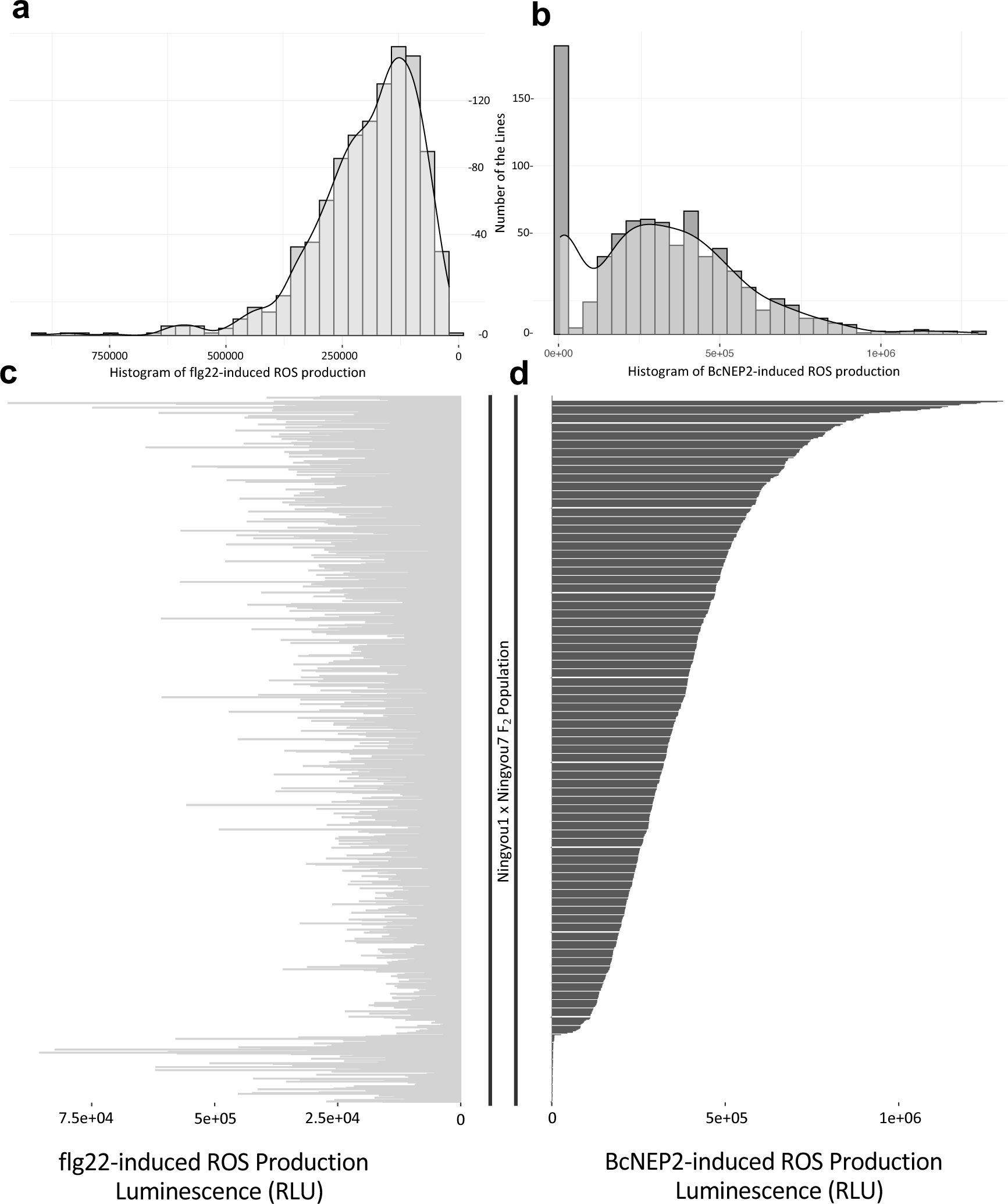
Phenotype data of each of 925 individuals from the Ning1×7 F_2_ *B. napus* population. The order of the individual lines from the population is arranged according to the increasing magnitude of the BcNEP2-induced ROS production. **(a)** Histogram and density plot of flg22-induced ROS production of the F_2_ population **(b)** Histogram and density plot of BcNEP2-induced ROS production of the F_2_ population **(c)** ROS production in response to 20nM of flg22. **(d)** ROS production in response to 50nM of BcNEP2. Data represent total RLU read over 40 minutes.

The recognition of the BcNEP2 molecule in this segregating population is binary (yes/no phenotype) **(Figure 2d)**, enabling selection of “Responsive” and “Non-responsive” pools. The level of NLP-induced ROS produced by responding individuals has a normal distribution **(Figure 2b),** enabling creation of BSA pools for “High responders” and “Low responders” potentially enabling the identification of genes modulating the NLP response.

### 3. BSA identifies variants highly associated with NLP-recognition located on chromosome A04

BSA was used to map binary and quantitative response for NLP recognition. DNA pools were generated with individual F_2_ plants selected from 5% of the extreme parts of the segregating population based on the binomial response to NLP (on/off), the magnitude of NLP response, and the magnitude of flg22 response. Based on the calculation of 5% of the 747 NLP-Responsive F_2_ plants, 30 individual F_2_ plants were sampled per pool. The details of the selected F_2_ individual with their corresponding 50nM BcNEP2 and 20nM flg22 ROS phenotype are shown in **Table 1**, **Figure S3**. Around ∼1.8billion raw sequence reads were obtained from each pool with a GC content of around ∼38% in all samples. This result is similar to that previously found with a GC content result of *B. napus* ZS11 genome with ∼40% GC content, with higher GC content in telomeric regions (Song et al., 2020).

After the quality control check, the reads were aligned to the *B. napus* cv Darmor-*bzh* reference genome (Chalhoub et al., 2014). To increase the coverage of the variations most highly associated with the responsiveness to NLP, “Responsive” (Pool3 and Pool4) and “Non-responsive” (Pool1 and Pool2) pools were merged *in silico*. In total 4,788,639 variations (SNPs and Indels) were found, of those 3,713,757 could be anchored to chromosomal positions on the Darmor-*bzh* genome. The quality of most of the variations in C02 and C09 was lower than the other chromosomes **(Figure S4)**. After excluding variants identical between the parental lines and those with coverage<20, and quality under 2,000, the total number of variants anchored to chromosomes decreased to 557,343. Each variant was scored with their corresponding Bulk Frequency Ratio (BFR) value (Trick et al., 2012).

A Manhattan plot was created with corresponding calculations of the BFRs of each variant on each specific chromosome is shown in **Figure 3a**. Peaks associated with NLP responsiveness were obtained on chromosomes A04, A09, C01, and C04. On chromosome A09, 3 variations with high BFR values were identified. Further evaluation of those variations showed that all 3 variants have a predicted effect on a *BnaA09g06090D* gene, a homologue of *A. thaliana myb96* (**Table S7**). MYB96 is a transcription factor involved in the activation of the genes that have a role against drought stress through abscisic acid (ABA) signalling (Seo and Park, 2010). On chromosome C01 there was only one peak associated with NLP responsiveness and the predicted effect of the associated variants were detected on the gene *BnaC01g28470D* **(Table S7)**. After close examination on this gene, it is found that the gene is annotated as 1803 bp long with 2 exons 247 bp and 39 bp long and there is no specific protein domain or *A. thaliana* homologue that can be assigned as related to this gene. A probable explanation for this is that the region is not well annotated on Darmor-*bzh* genome. Therefore, the valid data suggests that there is no predicted effect of this gene on the loss of NLP-recognition. The peak on C04 chromosome was evaluated further for effect predictions and functional annotations which identified a peak on a putative RLP homologue gene BnaC04g42570D **(Table S7)**. However, it has been previously shown that from 6 species within the Brassica clade only the species containing A or B genome are capable of recognising NLPs derived from Brassica pathogens (Schoonbeek et al., 2022). Therefore, the focus for further work was to define the candidate receptor gene located on chromosome A04.

**Figure 3.**
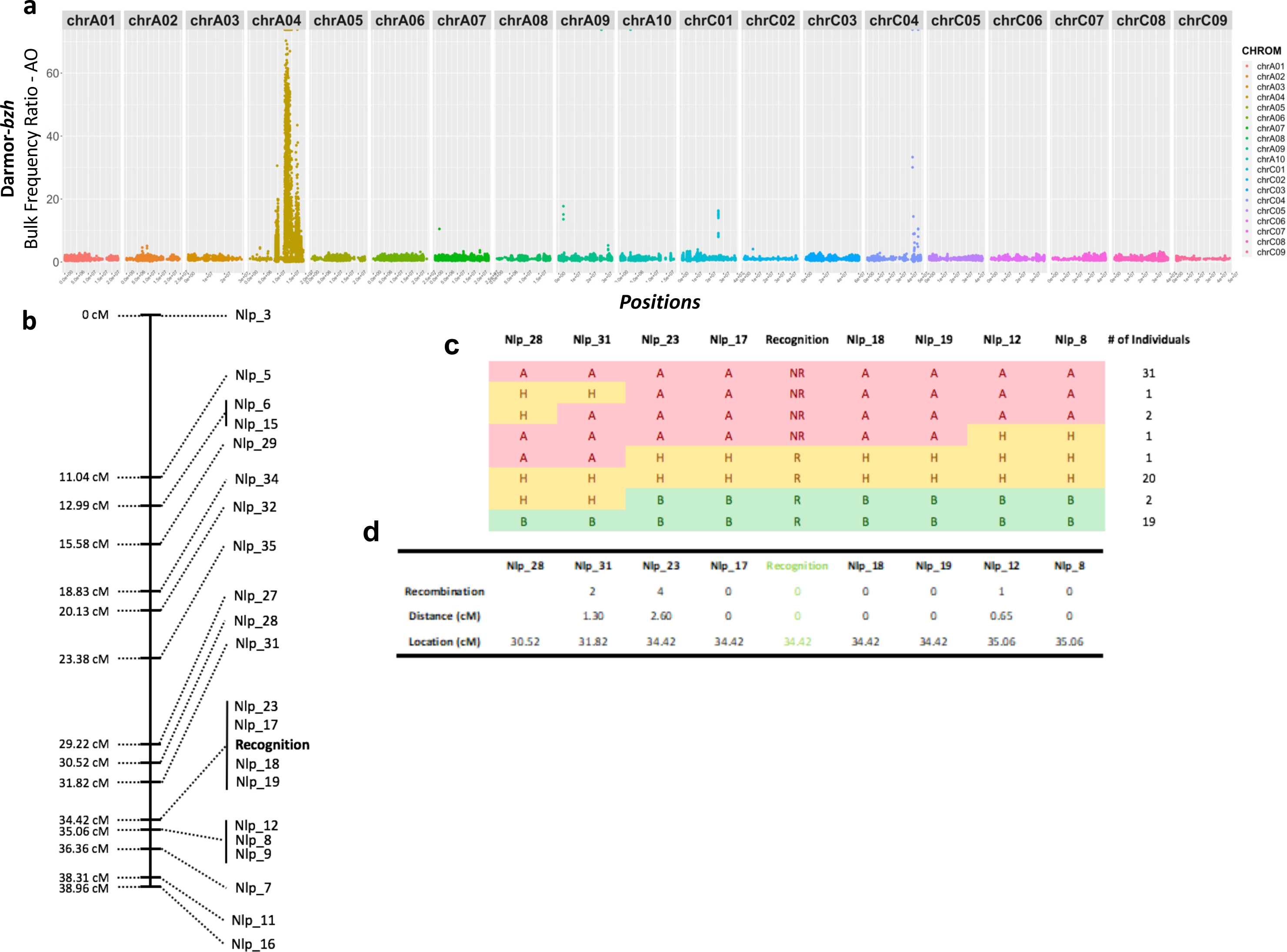
Genetic map of the NLP responsive region of the Darmor-*bzh* genome highly associated with “BcNEP2-recognition”. **(a)** Manhattan plot of the Bulk Frequency Ratios of variants called through *in-silico* merged “Responsive” and “Non-responsive” pools. After excluding the variants with lower than 20 read depth, 2000 quality and the same between parents, the positions of the 557,343 variants are plotted along X-axis on full genome. **(b)** QTL map of “BcNEP2-recognition” region of 77 plants from F_2_ plants with 21 KASP markers. **(c)** Genotypic features of each individual in the testing panel (in total 77 plants) for 8 KASP markers that are closely linked to BcNEP2-recognition (NR; Non-responsive (Red), R; Responsive (Green), H; Heterozygous (Yellow)). **(d)** Details of each marker; number of recombinations between the adjacent markers and their corresponding locations on the generated genetic map.

The variants with highest BFR are located on the A04 reaching up to 70 **(Figure 4c)**, supporting the results from the AT **(Figure 1)**. Variants with BFR value of infinity were also identified showing presence/absence polymorphisms between bulks. After applying the filters described above, the number of variants on A04 was detected as 24,660 **(Figure 4c)**. On A04, 4 main peaks were obtained, the genomic length of the 1st and the 3rd peak was ∼1 Mbp, the genomic length of the 4th peak was ∼200 kbp and the genomic area covered by the highest peak (2nd) was spread on ∼ 3 Mbp.

**Figure 4.**
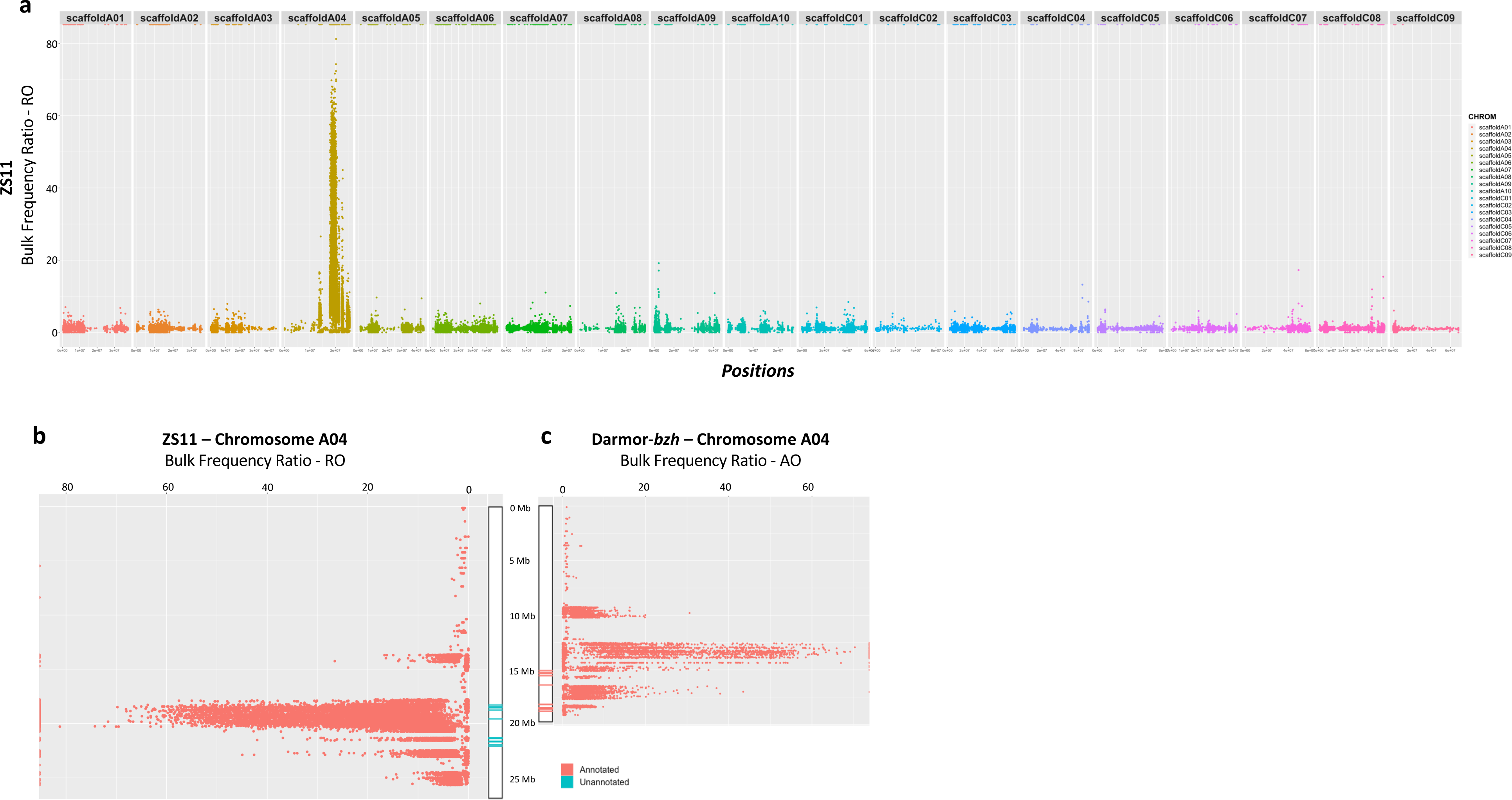
Manhattan plots of the Bulk Frequency Ratios of the variants called through *in-silico* merged “Responsive” and “Non-responsive” pools on the ZS11 genome. **(a)** After excluding the variants with lower than 25 read depth, 1500 quality and the same between parents, the positions of the 801,366 variants are plotted along X-axis on the ZS11 genome. **(b)** The positions of the 37,362 variants plotted along the X-axis on ZS11 A04 and **(c)** The positions of the 24,660 variants are plotted along the X-axis on Darmor-*bzh* A04. Regions containing homologues of *AtRLPs* which are annotated to a *Bna* gene in Darmor-*bzh* are illustrated in “Orange”, related regions present in ZS11 but not annotated to any *BnaZS* gene are illustrated in “Blue”.

### 4. Fine mapping of the region associated with NLP-recognition using KASP Markers

To construct a genetic map and delimit the region associated with the NLP responsiveness, KASP markers **(Table S2)** were designed from highly associated SNPs identified in the BSA analysis. Two key features of the SNPs were used as criteria for deciding the candidate SNPs to design KASP markers. The computed BFR AO value (Bulk Frequency Ratio from alternative allele) was the first feature as the values selected to be between 40 and 72. Second, the functional predicted effects on the annotated genes of the SNPs **Table S8**. The list of the SNPs used in KASP marker design, along with their corresponding features, is shown in **Table S9.**

A DNA test panel with samples from 77 F_2_ plants and parental lines was used to validate the 41 markers with specific FAM tail and HEX tail. The test panel included 6 lines that were responsive to NLPs to confirm the marker is diagnostic on lines external to the mapping population. 21 out of 41 markers were suitable for genetic map construction with their p-value <1.0E^-10^. Also, nine out of 21 KASP markers were able to validate the responsiveness in the other 6 lines out of the F_2_ BSA population **(Figure 3b)**. A quantitative trait locus (QTL) locus for a hypothetical receptor gene was mapped on the A04, and the SNPs highly associated with the NLP response phenotype was delimited to a maximum of 3.25 cM. The flanking markers NLP_31 and NLP_12 are 2.6 cM and 0.65 cM away from the markers that are linked to the locus of interest respectively **(Figure 3c)**. There are four markers (NLP_23, 17, 18, and 19) that co-segregate with NLP recognition in this population **(Figure 3d)**.

### 5. Comparative Coding Sequence (CDS) search on two reference genomes at the region associated with NLP-recognition

For the most significant peak on A04, 245 annotated genes were found to be associated between marker BnDar.A04.12522928.C and BnDar.A04.17468291.C (**Table S10**). The number of associated genes detected via BSA analysis was therefore too many to identify a single candidate gene. Among genes highly associated with the trait, homologues of Arabidopsis RLPs and those with LRR type domain features were filtered and listed in **Table S11**. However, of the 22 homologues of *Atrlp23* detected on the Darmor-*bzh* genome (Schoonbeek et al., 2022), the four located on chromosome A04 did not show any significant differences between the bulks and so were not identified as likely candidates. Additional analysis was therefore required to reduce the list of candidate genes.

Although Darmor-*bzh* is the most improved and well-studied reference genome, it is non-responsive to NLPs (Schoonbeek et al., 2022), and therefore genes responsible for NLP response are likely to be absent or non-functional. In 2020, the genome sequence for Zhongshuang II (ZS11) was published (Song et al., 2020), and ZS11 is NLP-responsive.

To further reduce the region associated with NLP-recognition and to find possible candidate NLP receptor genes, the genomic data from bulks were aligned to ZS11 reference genome. The BSA pipeline was performed and in total 3,672,282 variants (SNPs and Indels) were found as anchored to ZS11 chromosomes. To exclude the false positives from the analysis, the thresholds were set at 1500 for quality and 25 for depth for the variants. After filtering steps, the total number of variants were 801,366 (**Figure 4a)**. From them 37,362 of the variants were anchored to chromosome A04. As expected, the highest peak was identified on A04, suggesting a strong association to the trait (**Figure 4b**). Between marker Bn.ZS.A04.17867537.C and Bn.ZS.A04.22885838. A in total 405 annotated genes were found to be associated with the trait **(Table S12)**.

To further evaluate the region associated with NLP-recognition on ZS11 chromosome A04, Darmor-*bzh* genes were extracted and BLAST search was performed against chromosome A04 on the ZS11 genome. In total 23 RLP homolog genes were found at the associated region on Darmor-*bzh* genome. All of those were also detected on the ZS11 genome with more than 80% identity. However, not all RLP homolog genes available on Damor-*bzh* were annotated on the ZS11 reference genome (**Table S13**).

Since not all RLP homologs are completely annotated along the ZS11 genome (**Figure 4b**), RNA-Seq data was used to define the un-annotated genes across the associated region. The ZS11 RNA data were aligned to the reference genome and the presence of potential RLP homologs was investigated on A04. One potential candidate gene could be defined based on the presence/absence difference between the genomic data from the bulks and the parental lines. The un-annotated expressed putative gene located on ZS11 chromosome A04:20,675,496-20,678,125 was present in the NLP-responsive parent (Ningyou1) and absent in the non-responsive parent (Ningyou7) (**Figure S5**). This candidate region is located in the intronic region of *BnaA04G0210300ZS*. In addition, there are anomalies with that annotation, as the length of the gene in total is 23.7 kbp (found between position 20,673,147 and 20,696,865) and it includes two exons, one of 190 bp length and a second of 90 bp length. Moreover, three other unannotated RLP homologs were located in this intronic region **(Figure S5)**. Alignment of RNA-Seq data further revealed that 10 out of 23 unannotated RLP homologs within the associated region are expressed, which supports the presence of unannotated putative genes **(Table S13)**.

### 6. BcNEP2-recognition enhances resistance to *B. cinerea* in *B. napus*

The Ning1×7 population was used to investigate the effect of BcNEP2-recognition on *B. cinerea* disease resistance in *B. napus*. There was a significant increase (p= 4.262e-07***) in resistance to *B. cinerea* in the BcNEP2-responsive lines which had smaller lesions compared to non-responsive lines **(Figure 5c, 5d)**. Whilst there is variation between the leaf samples taken from the same plant **(Figure 5e, 5f)**, the statistical differences in lesion sizes between the Responder and Non-responder groups were highly significant. The mean ± SD values of the lesion size for the *B. cinerea* infection of the Responder and Non-responder groups are 14.9 ± 2.8 mm and 18.3 ± 2.2 mm, respectively **(Figure 5g)**.

**Figure 5.**
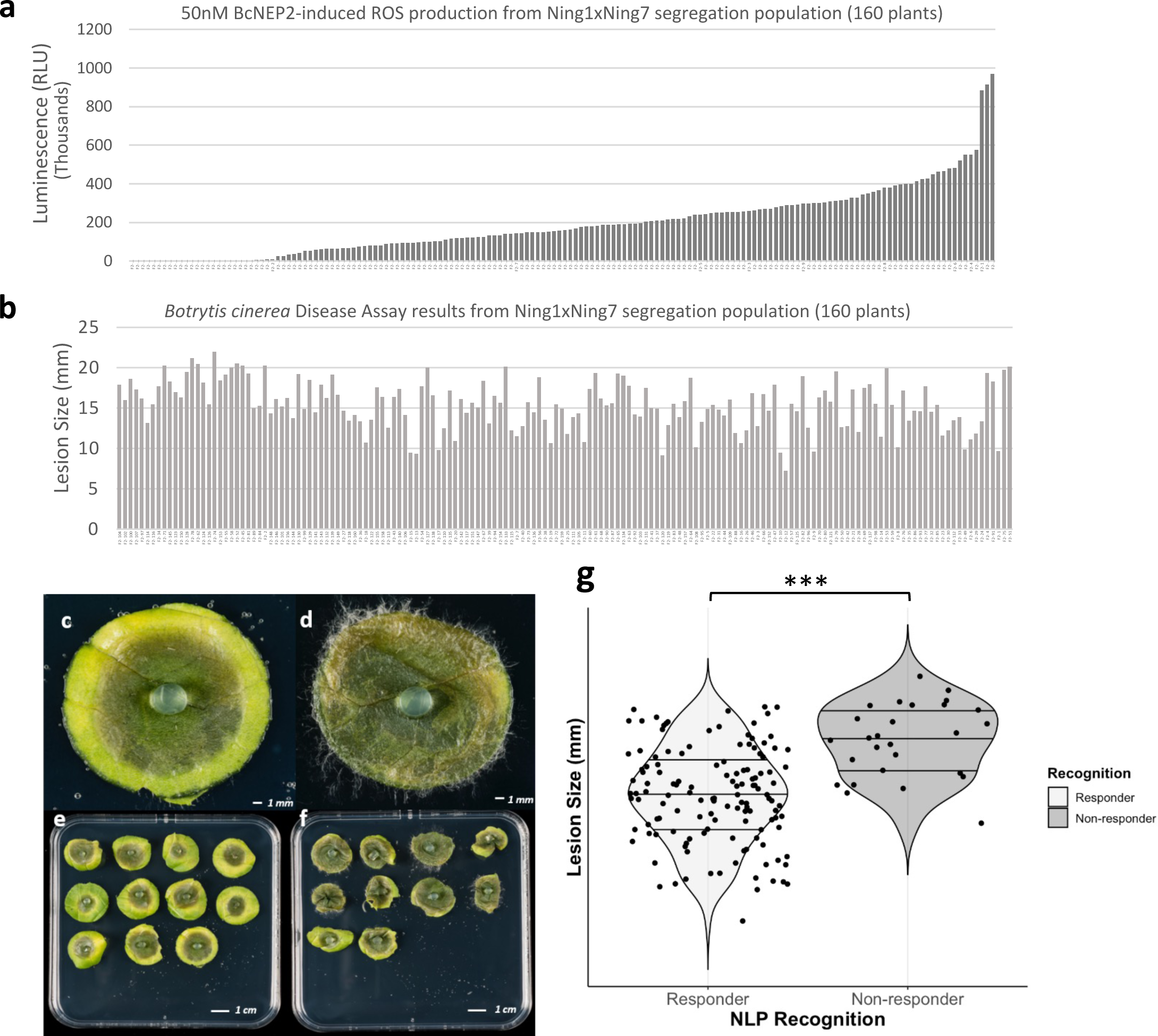
Phenotypic data of each of 160 individuals from Ning1×7 F_2_ population from BcNEP2-induced ROS production and *B. cinerea* disease assay. The order of the individual lines from the population is arranged according to the increasing magnitude of the BcNEP2-induced ROS production in the graphs. **(a)** ROS production in response to 50nM of BcNEP2. Data represent total RLU read over 40 minutes. **(b)** Lesion sizes of *B. cinerea* infection. Data represented as lesions size (mm) of *B. cinerea* infection on *B. napus* plant leaves 3dpi. **(c)** *B. cinerea* infected leaf disc taken from a “Responder” F_2_ plant. **(d)** *B. cinerea* infected leaf disc taken from Ningyou7 (Non-responder to NLP) which is the parental line of the F_2_ population. **(e)** A plate overview containing leaf disc samples from the “Responder” F_2_ plant. **(f)** A plate overview containing leaf disc samples from the “Non-responder” F_2_ plant. Five-week-old *B. napus* leaf discs were infected with *B. cinerea* plugs. Leaves were photographed at 3dpi. **(g)** Violin plot illustrating the distributions of lesion sizes belongs to 2 different groups of 133 Responder (light gray) and 27 Non-responder (dark gray) F_2_ individuals, showing the significant differences between the groups (p-value = 4.262e-07***).

To investigate whether the magnitude of MAMP-induced ROS production influences disease resistance, *B. cinerea* lesion sizes of 60 individual plants, classified as low and high responders to each MAMP, flg22 and BcNEP2, were compared. There was no significant difference between ‘low’ and ‘high’ responders of BcNEP2 and flg22 for *B. cinerea* disease resistance. (p-value_flg22_ = 0.597, p-value_BcNEP2_ = 0.439). For flg22, the shape of the plots was highly similar for both ‘low’ and ‘high’ BcNEP2-responders. **(Figure S6a, b)**. The mean ± SD value of the lesion size for the low BcNEP2-responders is found as 15.3 ± 2.7 mm which had more plants with bigger lesion sizes; however, with 14.7 ± 3.2 mm lesion size, high BcNEP2-responders were not significantly different. Overall, the resistance of *B. napus* to *B. cinerea* infection is significantly affected by the ability to respond to BcNEP2, but not by the magnitude of either BcNEP2 or flg22-induced ROS response.

### 7. *B. napus* plants non-responsive to BcNEP2 have significantly higher flg22-induced ROS production

Interestingly, the ability of the plant to respond to BcNEP2 affects the magnitude of the response to flg22. Although the flg22-induced ROS responses of BcNEP2 responders and non-responders’ was similar for the 1^st^ quartile value (40,455 RLU and 51,123 RLU respectively), the median values were were quite distinct from each other (Median_Responder_ = 61,438 RLU, Median_Non-responder_ = 100,187 RLU). There was a significant difference between the BcNEP2-responsive and non-responsive F_2_ plants in their flg22-induced ROS response phenotype with the mean ± SEM value as 135,349 ± 20,872 RLU and 82,985 ± 5,700 RLU respectively (p-value=0.00247 **) **(Figure 6a)**. When only the BcNEP2-responsive F_2_ plants were used in the correlation analysis, the results showed that there is a strong positive correlation between the magnitude of ROS production in response to BcNEP2 and flg22 with a correlation coefficient of 0.58 **(Figure 6b)**.

**Figure 6.**
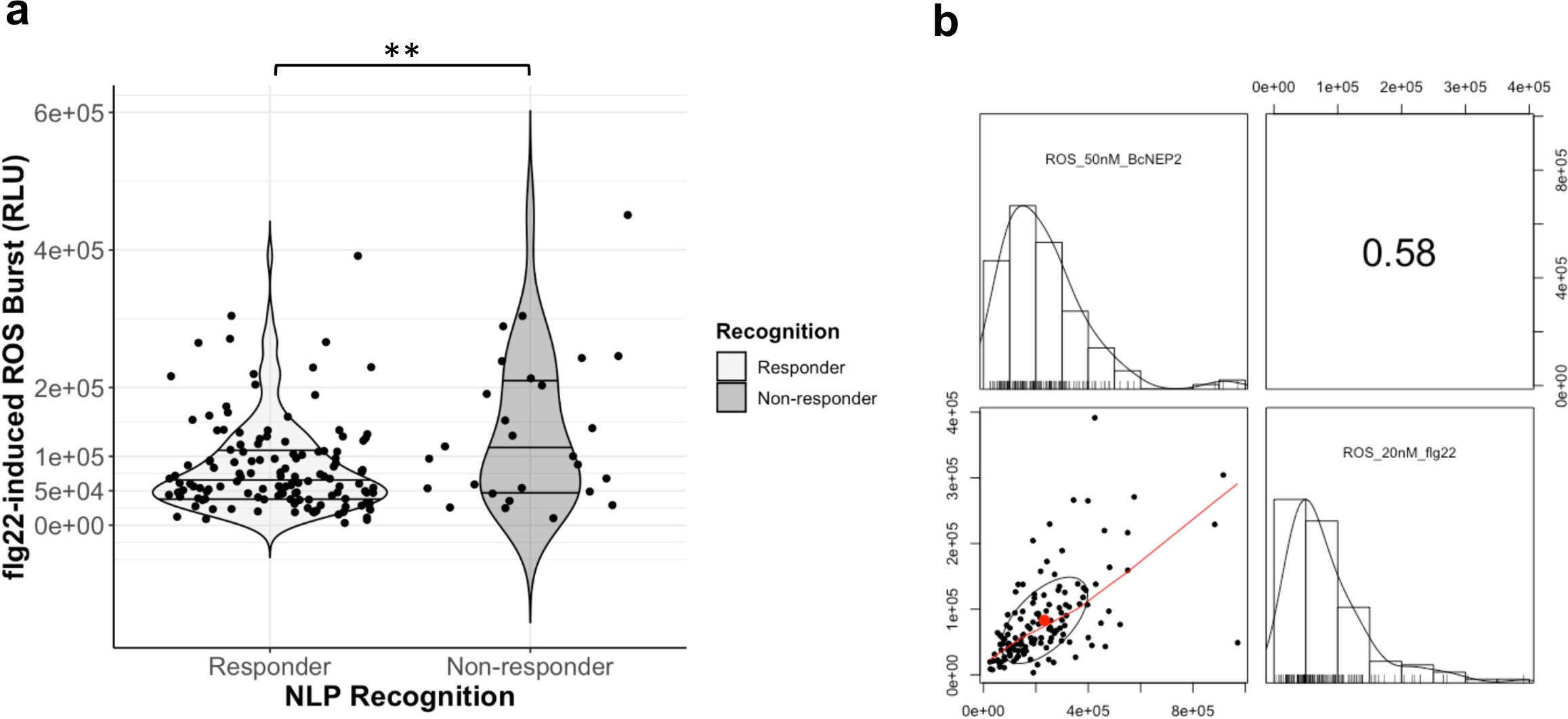
BcNEP2-recognition affects the magnitude of flg22-induced ROS production. **(a)** Violin plot illustrating the distributions of flg22-induced ROS production belongs to 2 different groups of 133 Responder (light gray) and 27 Non-responder (dark gray) F_2_ plants, showing the significant differences between the groups (p = 0.00247 **). **(b)** Correlation analysis between the 50nM BcNEP2-induced ROS production results of individual F_2_ plants from the Ning1×7 population and corresponding 20nM flg22-induced ROS production results. Histogram plots integrated with rug plots of total RLU in response to 50nM BcNEP2 (upper left) and 20nM flg22 (lower right), respectively. The scatter plot (lower left) was drawn with LOESS smooth showing the positive correlation between the results. The correlation coefficient value is in the upper right.

### 8. *BnaBSK1.A01* is significantly associated with the high BcNEP2-induced ROS production

The parallel phenotyping with BcNEP2 and flg22 enabled us to compare regions responsible for modulating the responses for each MAMP motif on *B. napus* genome.

Genomic data from Pool4 (High BcNEP2-responsive) and Pool3 (low BcNEP2-responsive) were run in the BSA pipeline to identify genes potentially modulating the magnitude of the BcNEP2-induced ROS production. In total, 3,801,929 variants were found as anchored to chromosomal positions on the *B. napus* genome, after filtering, 2,676,098 variants remained with more than 2000 quality score and 20 depth. After calculation of BFR values to identify variants most associated with the high BcNEP2-responsiveness, results clearly showed that the regions associated with the BcNEP2-recognition (**Figure 3a**) and the magnitude of BcNEP2-induced ROS production (**Figure S7a**) are different.

To detect the variants associated with high flg22-responsiveness, the same protocol was followed with Pool1 and Pool2 **(Table 1)** which are significantly different from each other in magnitude of flg22-induced ROS production. The highest association were located on A04 with a BFR value more than 20, which was similar to the result from high BcNEP2-responsiveness, but it is much more defined and clearer for High flg22-induced ROS response. Additionally, a peak at the beginning of the A02 was identified which is only associated with high flg22-induced ROS production (**Figure S7b**). The peak is mostly spread on ∼200 kbp, between position 300 kbp and 500 kbp. In that region, 28 annotated genes were found to be likely affected with the highest BFR value of 13.9 **(Table S14)**. Another peak located on A01; had a higher association with high BcNEP2-induced ROS production when compared with flg22-induced ROS production. Annotation and Effect prediction of 106 variants on A01, associated with the high BcNEP2-responsiveness phenotype, were found to have predicted effect on *BnaA01g02190D* gene **(Table S15)**. This gene is a homologue of *Atbsk1* (*At4g35230*), which is *BR-SIGNALLING KINASE1* physiologically associated with *FLS2* (Shi et al., 2013).

### 9. Arabidopsis *bsk1-1* mutants have significantly lower BcNEP2-induced ROS burst and decreased resistance to *B. cinerea* infection

*Atbsk1* (*At4g35230*), *BR-SIGNALLING KINASE1*, physiologically associates with *FLS2* and has a major role in flg22 recognition (Shi et al., 2013) but its role in NLP recognition is unknown. To functionally test the role of BSK1 in BcNEP2-induced ROS production, *A. thaliana bsk1-1* mutants were used. Before screening, the *A. thaliana* mutant *bsk1-1* and the Col-0 were genotyped to confirm the mutations **(Figure S8)**.

To test the possible effect of a mutation on *BSK1* on BcNEP2-induced ROS production, *bsk1-1* mutant seeds and the Col-0 were compared in the ROS assay using different concentrations of BcNEP2 and flg22. There was a gradual decrease in ROS production corresponding to decreasing concentration of both MAMPs. Similar results were observed in Col-0 by both BcNEP2 and flg22 MAMP motives. In contrast to the results of Shi et al. (2013), there was no significant difference between the wild type and *bsk1-1* mutant line for any concentration of flg22. However, unlike the flg22 response, a significant difference was found between the *bsk1-1* mutant and the Col-0 in response to 50 nM BcNEP2. Also, there was a clear trend of the BcNEP2-response of *bsk1-1*, being reduced compared to Col-0 at all concentrations. This significant difference shows that BSK1 is involved in BcNEP2-triggered ROS production in Arabidopsis **(Figure 7a, 7b)**.

**Figure 7.**
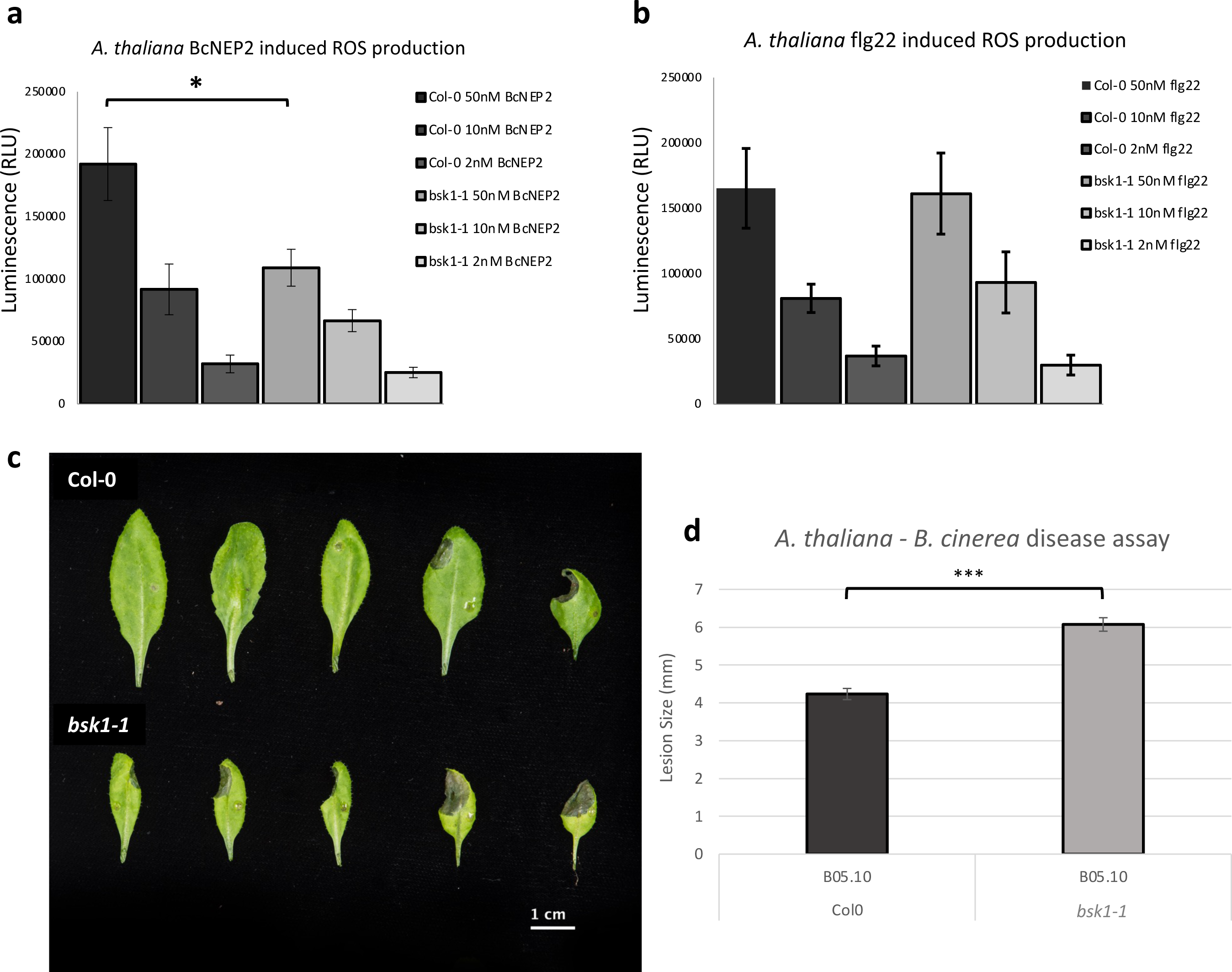
BSK1 is involved in BcNEP2-induced ROS production and *B. cinera* disease resistance. Leaves of the Col0 (wild type), *bsk1-1* mutant lines were treated with (a) 50nM, 10nM, 2nM BcNEP2 and (b) 50nM, 10nM, 2nM flg22 incubated with luminol and horseradish peroxidase to detect ROS. Data represent total RLU read over 40 minutes. Error bars are the standard error of 8 biological replicates. (p-value=0.0287*). *B. cinerea* disease screening of leaves from Arabidopsis plants. (c) Five-week-old Arabidopsis plants were infected with *B. cinerea*. 5μl drops from B05.10 strain placed on the upper part of the leaf. The plants were photographed at 3dpi. (d) Data represented as lesion size of *B. cinerea* infection on Arabidopsis plant leaves 3dpi. Error bars represent standard error of 12 biological replicates (p = 1.12E-14***).

The involvement of co-receptor BSK1 in disease resistance against the *B. cinerea* was tested using Col-0 and *bsk1-1* mutant lines. The *bsk1-1* mutant line showed significantly higher susceptibility to *B. cinerea* wild type strain with larger lesions compared to Col-0 (**Figure 7c, 7d**).

### 10. Defence-related genes are differentially induced in Col-0 and *bsk1-1* mutant line with BcNEP2 pre-treatment

The induction patterns of the genes having a key role in the JA pathway such as *JAR1* (Suza and Staswick, 2008) and a transcription factor involved in plant defence, *WRKY33* (Birkenbihl et al., 2012; Zheng et al., 2006) were examined after BcNEP2 treatment. Also *BSK1* and *RLP23* gene expressions were investigated using RT-qPCR. The *bsk1-1* mutation neither negatively nor positively affects the expression levels of the *BSK1* and *RLP23* genes. Additionally, there was no significant induction due to BcNEP2 treatment for these 2 genes when compared with the mock-treated samples **(Figure S9)**. There was significant induction in *WRKY33* expression of the wild type BcNEP2-treated plants. In contrast, the *bsk1-1* mutant line did not show any significant induction of *WRKY33* at the same time points when compared with its mock-treated plants. There was a trend of increased induction at 12hpi (hours post-treatment) in the mutant line, returning to its basal level at 24hpi **(Figure 8a, 8b)**. A similar result was observed for *JAR1* gene induction. At 12hpi, there was significant induction in *JAR1* expression in both wild type and mutant lines, whereas, at 24hpi, the significant induction was only detected in wildtype BcNEP2 treated plants when compared with the mock-treated ones **(Figure 8c, 8d)**.

**Figure 8.**
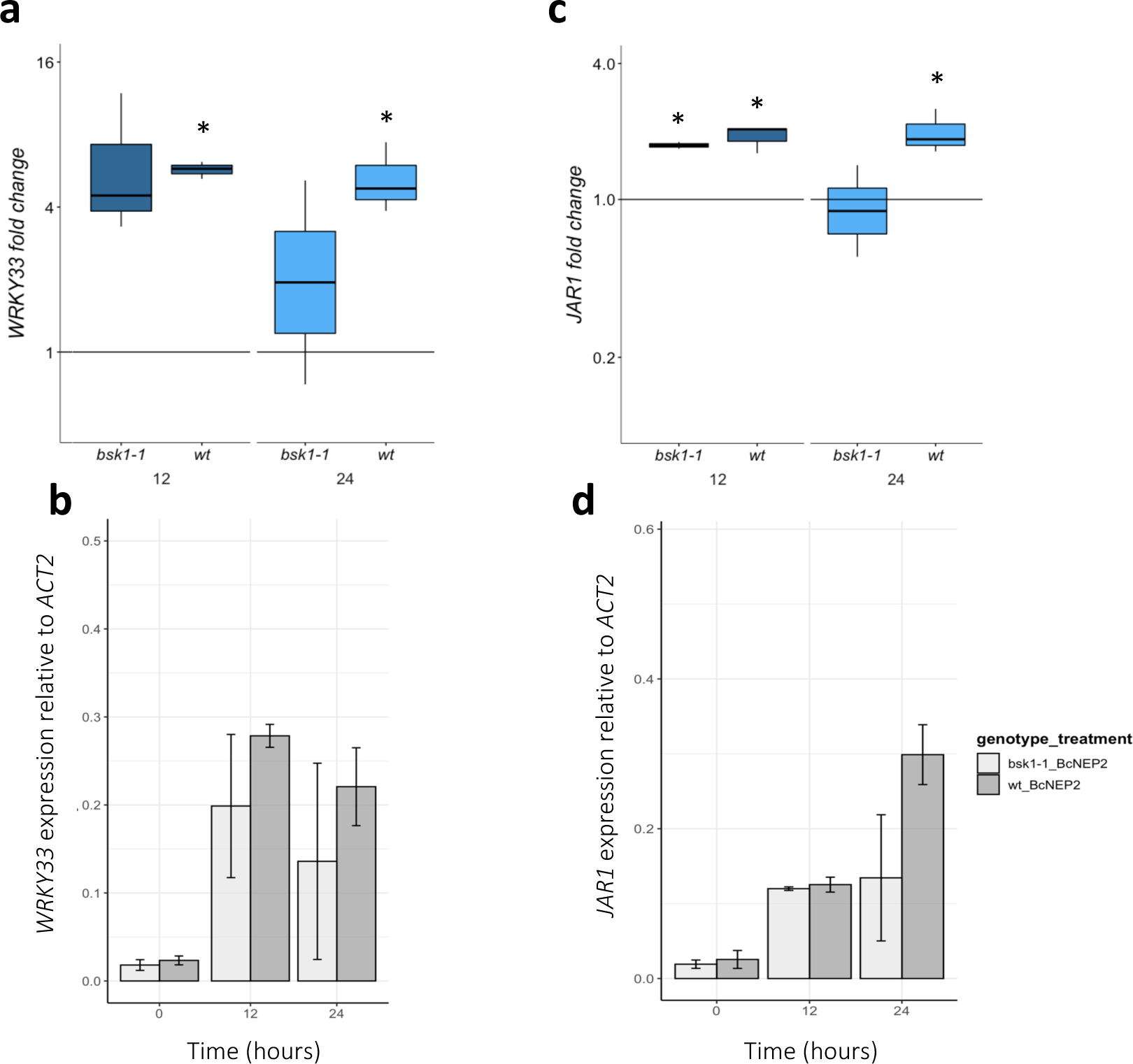
Quantification of induction and expression of *WRKY33* (a,b) and *JAR1* (c,d) genes of Arabidopsis after 100nM BcNEP2 treatment at 12 and 24 hours post treatment. mRNA levels of the genes were measured by qRT-PCR in BcNEP2-treated and mock-treated Arabidopsis plants. *ACTIN2* was used as a normaliser. Bars represent means of three biological replicates ± SEM. Asterisks indicate significant differences between expression levels in mock and treated leaves (Student’s t-test *, P <0.05).

## Discussion

Our results demonstrate for the first time that NLP recognition is correlated with enhanced resistance to *B. cinerea* in *B. napus*. The result confirms the previous observation of RLP23-mediated resistance in Arabidopsis (Ono et al., 2020) but contrasts with the previous study in *B. napus* where no correlation was observed between NLP recognition and disease resistance (Schoonbeek et al., 2022). In that study two responsive *B. napus* cultivars (N02D, Ningyou1) were compared with non-responder cultivars (Tapidor, Ningyou7) in disease resistance against *B. cinerea* and no statistical difference was detected (Schoonbeek et al., 2022). However, it was also suggested that cultivars used in that study had quite different genetic backgrounds and there might be other factors affecting the disease resistance which could mask the effect of NLP-triggered immunity against *B. cinerea* disease resistance in *B. napus*. The considerable amount of genetic similarity between Ningyou1 and Ningyou7 parental lines (X. Wang et al., 2017, J. Zou et al., 2019) was effective at eliminating background genetic variation which might interfere with the disease assay results. This is the first demonstration that NLP recognition affects disease resistance in *B. napus*.

To map the genomic locus for BcNEP2-recognition a combination of AT and BSA approaches were used. AT identified two loci on A04 and C05 whereas the BSA confirmed only the A04 location. One explanation for this could relate to the different populations used in the analyses; AT uses transcriptome sequences from many genetically divergent *B. napus* accessions whereas the BSA will only contain the genetic variation present between the two parents used to develop the segregating population. Further variation controlling responsiveness, for example at the C05 locus, may be present within the other responding lines. Although Darmor is a non-NLP-responsive cultivar, it is the most widely used and well-annotated *B. napus* reference genome (Chalhoub et al., 2014). Using Darmor-*bzh* in BSA resulted in a well-defined and clear peak covering a ∼ 3Mbp region on A04. The region was delimited to a maximum of 3.25cM which could be covered with 21 KASP markers, four of which were co-segregating with NLP-recognition. Another gene in the region, *BnaA04g18600D*, was highly associated with the NLP-recognition with BFR value reaching up to 26 **(Table S16)**. *BnaA04g18600D* is a homologue of SOBIR1 which has a key role in NLP-recognition in Arabidopsis. SOBIR1 forms a complex with RLP23 and this functional complex then recruits BAK1 to start the signalling cascade during NLP recognition (Albert et al., 2015). The most likely receptor candidates are expected to be RLP-like genes since these are associated with the perception of the NLPs and initiating the signalling mechanism. Since Darmor-*bzh* is non-responsive to NLPs (Schoonbeek et al., 2022), genes responsible for NLP-recognition are likely to be absent or non-functional on this reference genome. Therefore, NLP-responsive Zhongshuang II (ZS11) reference genome (Song et al., 2020) was also used in this study. The BSA results confirmed the association on chromosome A04 with a clearly defined peak and enabled us to define the region to 405 annotated genes on the ZS11 genome.

A comparison of NLP-recognition-associated regions in *Darmor-bzh* and ZS11 revealed that, although there is a deviation in sequence similarity between the genomes, most of the genes detected were present on both genomes. However, not all RLP homologs from Damor-*bzh* were present in the current ZS11 gene annotation. Assuming that there are potential genes in the region that are not annotated on ZS11 yet, we used ZS11 RNA-Seq data to show that 10 out of the 23 RLP homologs from Darmor-*bzh* are expressed and located in the corresponding region of ZS11. This result demonstrates that there are unannotated putative RLP homologs in this region of the ZS11 genome. Despite the rapid advancement of sequencing technologies, gene annotation remains challenging, especially for large gene families. By the very nature of crop genetics (hybridization, etc), current annotations don’t capture the full diversity and although contemporary methods typically give a very good general overview of gene content, careful manual curation of target loci remains mandatory. Importantly, as illustrated in our example, gene comparisons based on annotations derived from different methods require a high level of scrutiny.

Our results provide the first indication that BSK1 could modulate BcNEP2-induced ROS production, possibly through its involvement in a complex similar to that reported for BAK1/SOBIR1 (Albert et al., 2015). Although the BcNEP2-induced ROS production was significantly reduced in *bsk1-1* mutant lines, there was no reduction in flg22-mediated ROS burst as previously reported (Shi et al., 2013). One possible reason for this could be the environmental conditions used in these experiments were different from those used by Shi et al. Since BSK1 is involved with BR signalling, it is reasonable to suggest that the environment could affect the result. BR sensing is involved in both the regulation of immunity and also of growth (Lozano-Durán and Zipfel, 2015), and it is also well-known that different results are obtained for Arabidopsis between labs (Massonnet et al., 2010). Altogether, these results suggest that BcNEP2-induced ROS production is affected to a greater extent in *bsk1-1* mutants than flg22-induced ROS burst. Additionally, to examine the molecular basis underlying the effect of *bsk1-1* mutation on BcNEP2-recognition, transcript abundance of genes from Ethylene (ET)- and Jasmonic Acid (JA)-mediated defence pathways that play a key role in immunity against *B. cinerea* in Arabidopsis (Birkenbihl et al., 2012; Staswick et al., 2002; Zheng et al., 2006), were investigated. The results suggest the involvement of BcNEP2-triggered PTI during the infection process of *B. cinerea* in Arabidopsis. In addition, the loss of *JAR1* induction at 24hpi of the bsk1-1 mutant might be due to a lack of induction in *WRKY33* gene expression, as it has been previously shown that this transcription factor prevents the antagonistic effect of SA-mediated pathways on JA-mediated responses (Birkenbihl et al., 2012).

Our results provide evidence of competition between PRRs that can affect the recognition of specific MAMPs. When compared to the BcNEP2-responsive group of plants, the BcNEP2-non-responsive group had a significant increase in the amount of flg22-induced ROS **(Figure 6a)**. In a previous study with Arabidopsis, the downstream immunity-related genes upregulated by cytotoxic NLPs strongly overlapped with the genes that are induced by flg22 (Oome et al., 2014; Oome and Van den Ackerveken, 2014; Wan et al., 2019) and the downstream components of signalling cascades are shared between the different PRRs (Albert et al., 2015). Thus, competition for the shared common co-receptors for different PRRs may operate during signal transduction, explaining the increased capacity to respond to the flg22 molecule in the absence of the NLP receptor. Receptor-Like Cytoplasmic Kinases (RLCKs) are common signalling modules (Liang and Zhou, 2018) and the expression of common co-receptors can be enhanced to intensify the immune responses gathered via multiple receptors (Kamoun et al., 2018).

## Conclusion

This study elucidates a series of investigations into the innate immune system of the *B.* napus and how it potentiates quantitative disease resistance of the plant. Our results show that BcNEP2-recognition induces resistance in *B. napus* and illustrates the value of translational research to transfer knowledge from model organisms to complex crop species. The genetic locus for NLP-recognition was mapped to a complex region on A04 containing RLP23 like homologues as well as other genes. Combined genomic and transcriptomic analysis identified several unannotated putative genes within the region as potential candidates for NLP-recognition on the ZS11 genome. Additionally, our results demonstrate that *BSK1* is involved in modulating the BcNEP2-induced ROS production and positively contributing to disease resistance against *B. cinerea*, linking results from the *B. napus* with mechanistic studies from Arabidopsis.

## Acknowledgement

HAY was supported by The Newton-Katip Celebi Fund given by the British Council and The Scientific and Technological Research Council of Turkey (TUBITAK). RHRG was supported by the UK Biotechnology and Biological Sciences Research Council (BBSRC) through the Designing Future Wheat (BB/P016855/1) Institute Strategic Program and also supported by the European Research Council (ERC-2019-COG-866328). CJR was supported by Biotechnology and Biological Sciences Research Council (BBSRC) grants BB/N005007/1 (MAQBAT) and the Plant Health Institute Strategic Programme BB/P012574/1 ‘Response’ subprogramme BBS/E/J/000PR9796. HJS and RK were supported by BB/N005007/1. RW, CNJ and GSS acknowledge financial support from Institute Strategic Programme ‘Genes in the environment’ (BB/P013511/1) and BBSRC Brassica Rapeseed And Vegetable Optimisation strategic Longer and Larger fund (BRAVO sLOLA) (BB/P003095/1). We thank the BRAVO sLoLa research team for access to gene expression data. We are grateful to Dingzhong Tang (CAS) for providing Arabidopsis *bsk1-1* mutant seeds. The authors would like to thank Horticultural Services in JIC for their valuable support in growing plants.

## Author Contributions

All in vivo experiments were conducted by H.A.Y with help from R.K. and E.V. who contributed to bulk sampling and phenotyping of the mapping population. R.W. and H-J.S. contributed to population development and experimental design. H-J.S. contributed to phenotypical data collection for AT analysis. H.A.Y. and R.H.R.G. performed all bioinformatics associated with the BSA, KASP marker design and mapping. C.N.J. performed AT analysis. B.S. and G.S.S. performed all bioinformatical analyses associated with RNA-Seq data. H.A.Y. wrote the manuscript. C.R. and R.W. supervised and co-directed the research and edited the manuscript. All authors read and approved the final manuscript.

## Supplementary Figures

**Figure S1.**
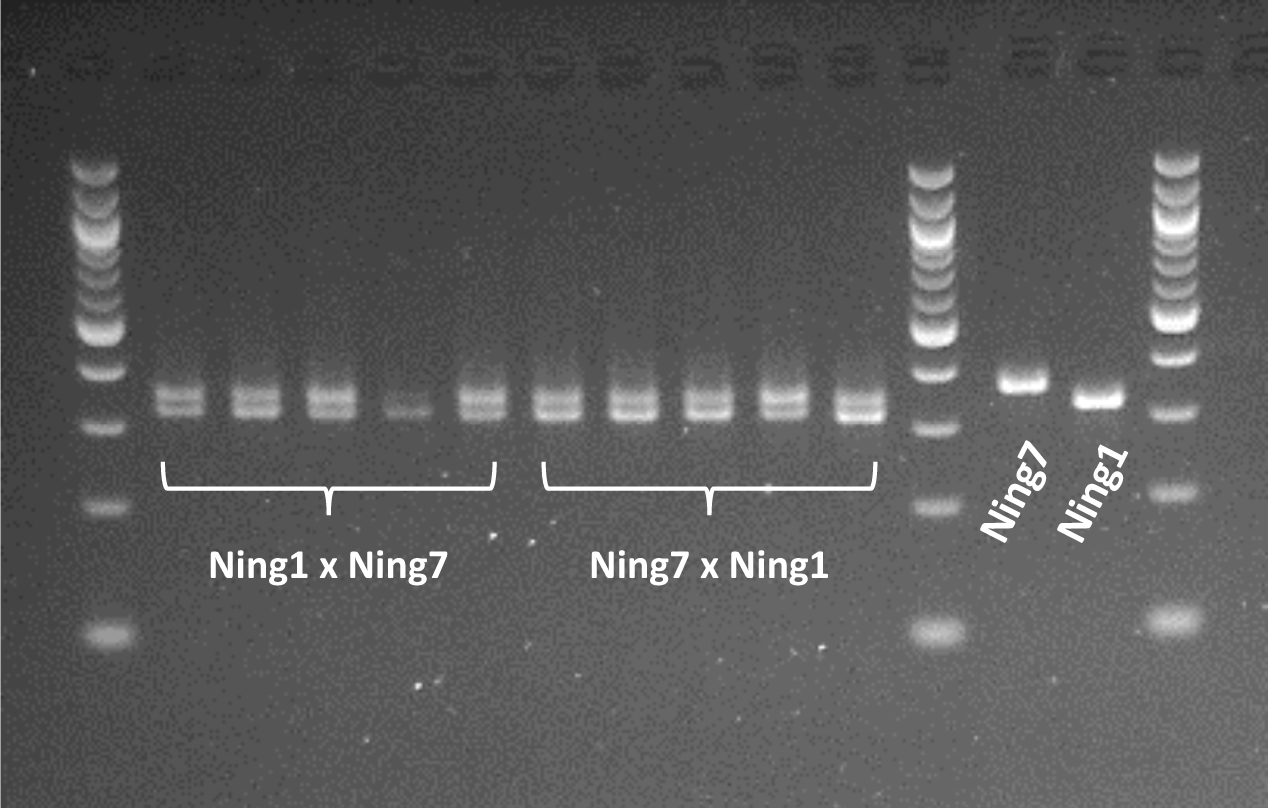
Amplification of microsatellite marker sR12095 in reciprocal crosses and parental lines. The fragments were separated in a 3.0% denatured Agarose gel. To visualize the amplified PCR products, 6× loading dye was added to PCR products before loading onto an agarose gel containing ethidium bromide and 15µl of PCR products were run on Agarose gel. The 1^st^, 12^th^ and 15^th^ lanes of are 100bp DNA Ladder (NEB #B7025). The rest of the lanes and their corresponding sample names are represented in the figure. Polymorphism of microsatellites among the Ningyou1 (Ning1) and Ningyou7 (Ning7) parents seen on the last lanes on the right side of the gel.

**Figure S2.**
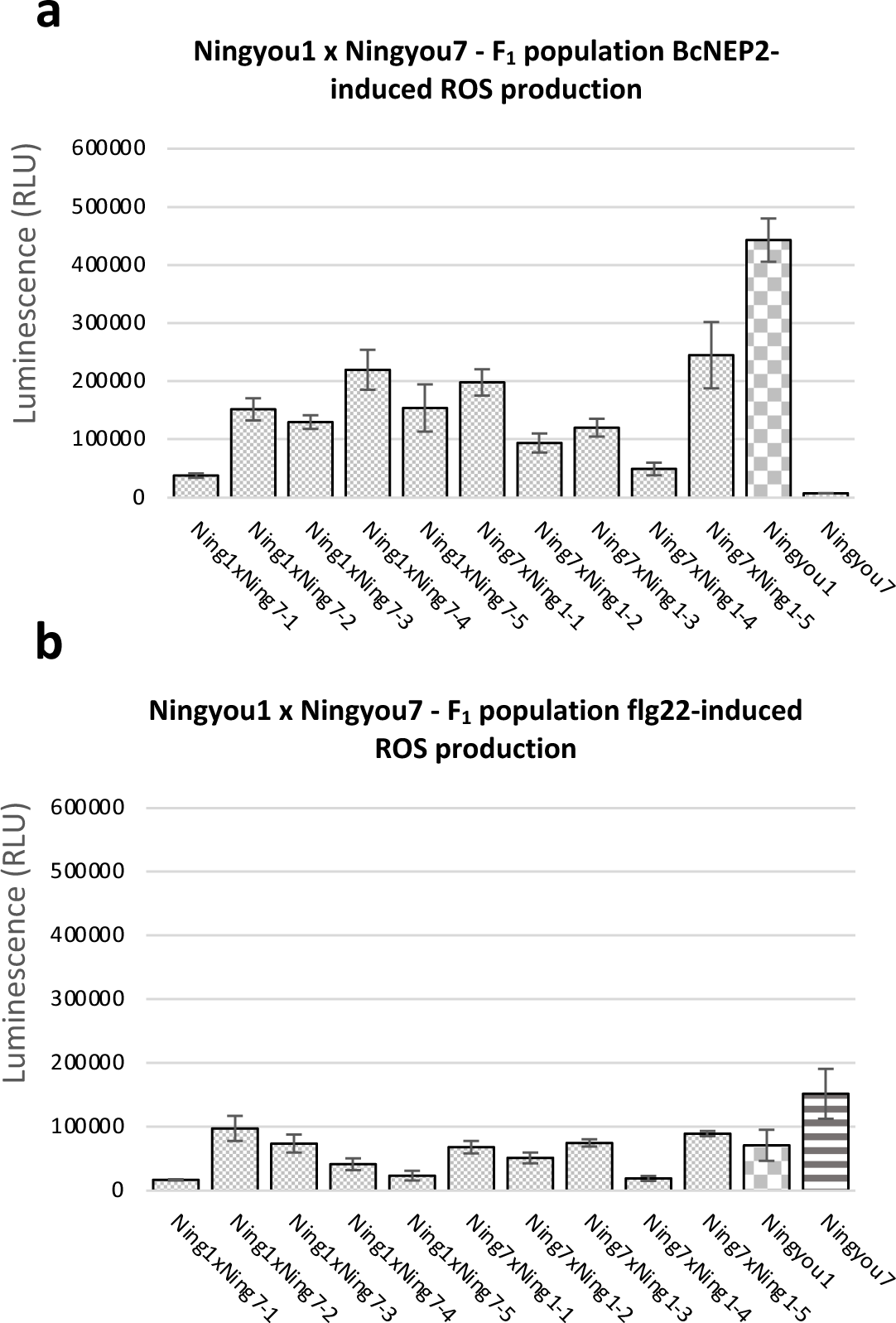
PAMP-induced ROS production from 10 F_1_ plants derived from 2 different reciprocal crosses **(a)** 50nM BcNEP2-induced ROS burst from 3^rd^ leaves of Ningyou1 x Ningyou7 F_1_ plants. **(b)** 20nM flg22-induced ROS burst from 3^rd^ leaves of Ningyou1 x Ningyou7 F_1_ plants. Bars represent Total Relative Luminescence Unit (RLU) read over 40 minutes. Error bars are the standard error of 8 biological replicates.

**Figure S3.**
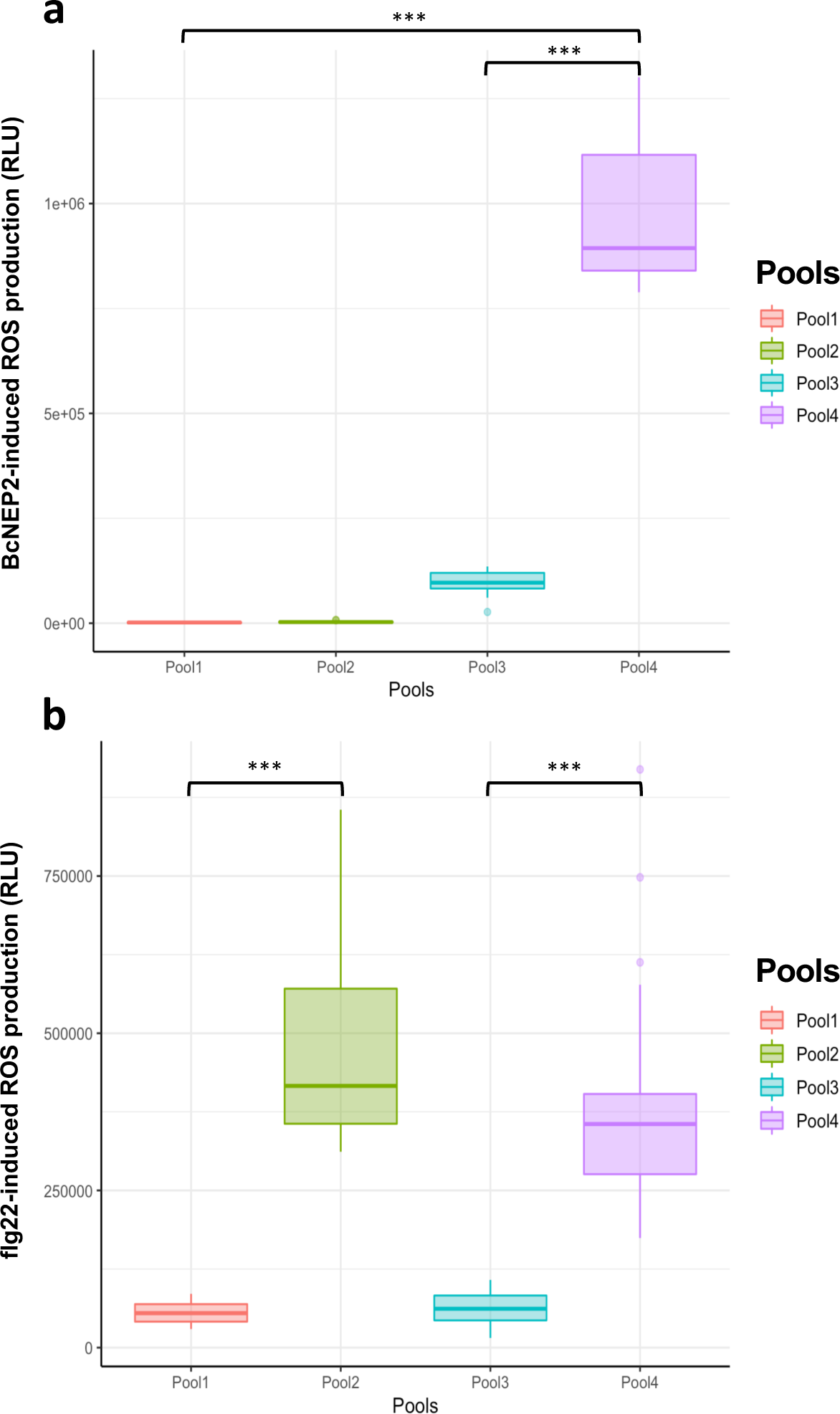
The difference between the pools based on their corresponding (a) 50nM BcNEP2-induced ROS production (b) 20nM flg22-induced ROS production. Box plot illustrating the distributions of ROS production belonging to 4 different pools of 30 F_2_ individuals from BSA population; Pool1 (Rose), Pool2 (Green), Pool3 (Blue), and Pool4 (Purple), showing maximum phenotypic differences between the pools. (***p-values are lower than 5.3e^-14^).

**Figure S4.**
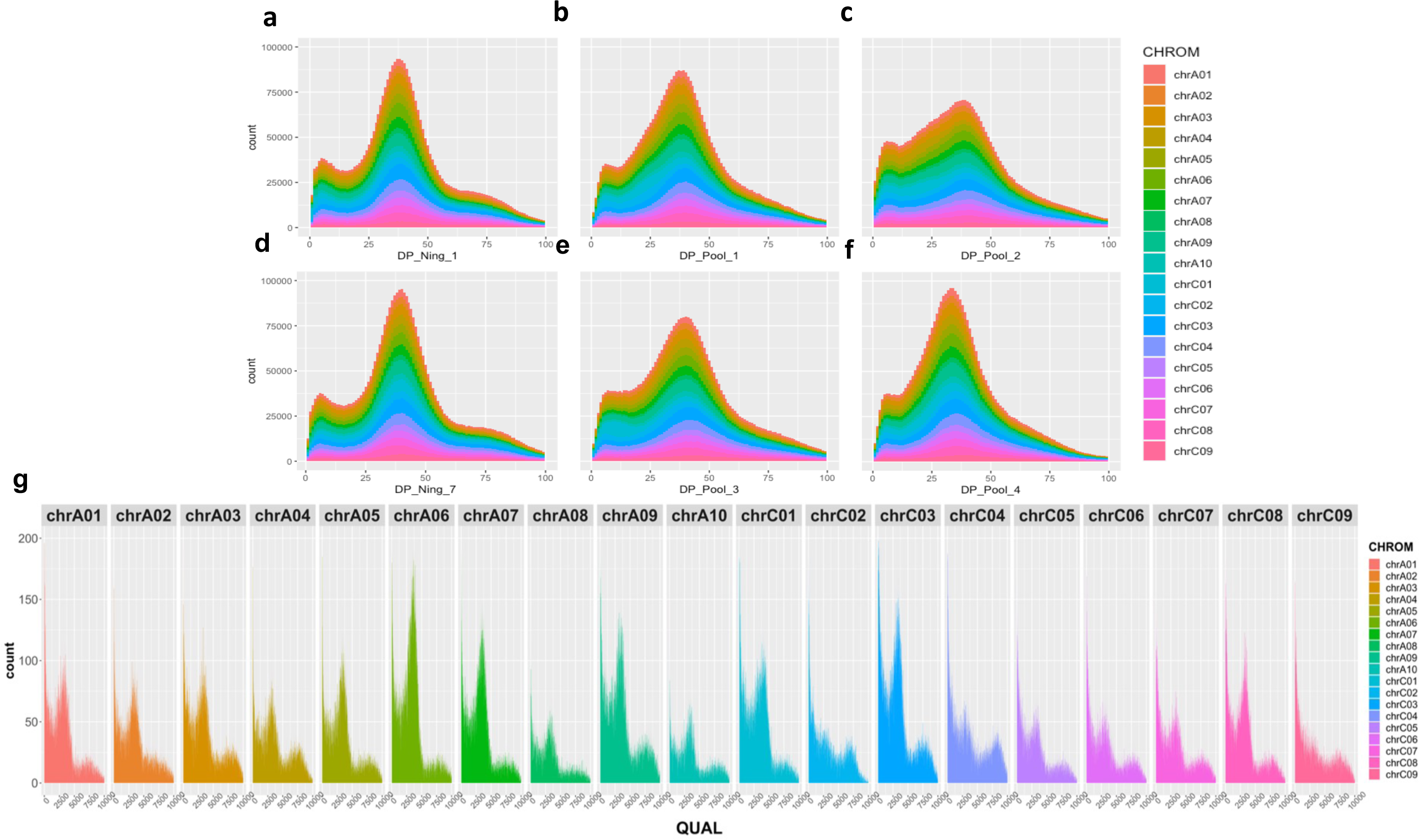
Histogram plot of depth (DP) and quality (QUAL) values for 3,801,929 variations colour coded based on their locations on the *B. napus* chromosomes. (bin width= 1). **(a)** Depth – Ningyou1 **(b)** Depth – Ningyou7 **(c)** Depth – Pool 1 **(d)** Depth – Pool 2 **(e)** Depth – Pool 3 **(f)** Depth – Pool 4 **(g)** Quality values of all variants in each corresponded chromosome (bin width= 1).

**Figure S5.**
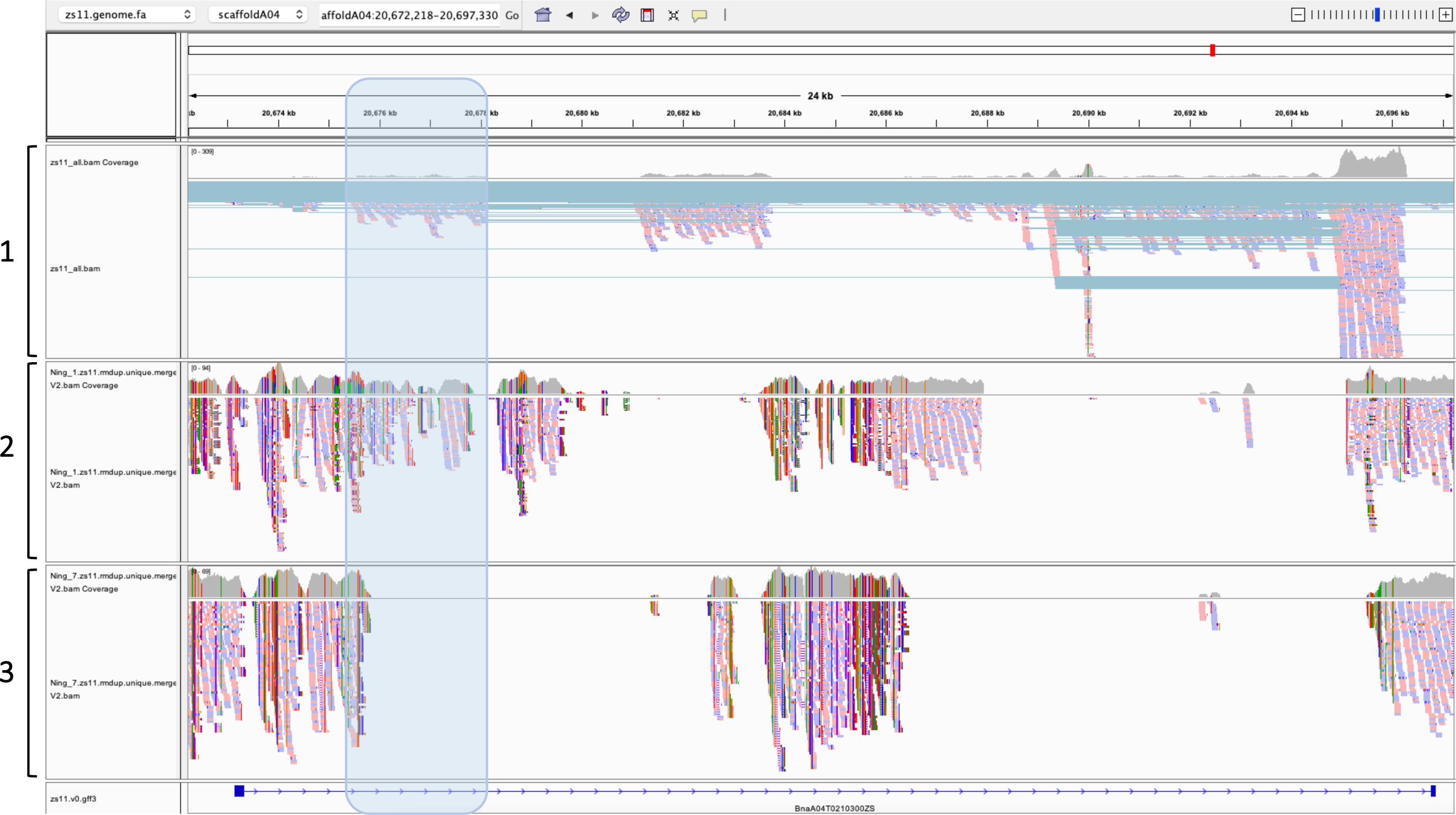
Illustration generated from Integrative Genomics Viewer (IGV) captured the area of ‘’scaffoldA04:20,672,218-20,697,330” on ZS11 genome. From top to bottom, the panels are representing the aligned data of (1) RNA-Seq data from ZS11, (2) DNA-Seq from Responsive parent (Ningyou1) (3) DNA-Seq from Non-responsive parent (Ningyou7). And the bottom panel representing the corresponded annotated gene *BnaA04G0210300ZS.* Region highlighted with Blue panel is captured the area of ‘’scaffoldA04:20,675,496-20,678,125” on ZS11 genome.

**Figure S6.**
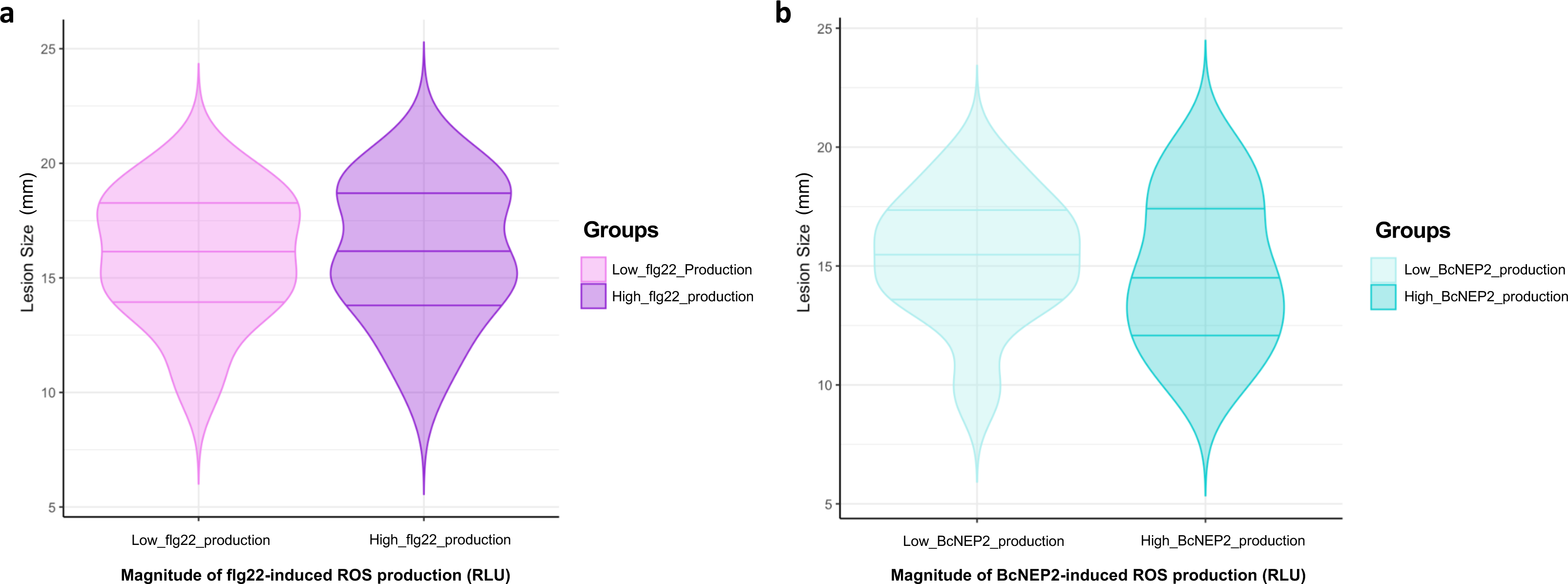
QDR against to *B. cinerea* infection is not affected by the magnitude of either BcNEP2 or flg22-induced ROS production. **(a)** Violin plot illustrating the distributions of lesion sizes belongs to 2 different groups of 30 Low flg22 Responder (Pink) and 30 High flg22 Responder F_2_ individuals (Purple). **(b)** Violin plot illustrating the distributions of lesion sizes belongs to 2 different groups of 30 Low BcNEP2 Responder (light blue) and 30 High BcNEP2 Responder F_2_ individuals (Blue).

**Figure S7.**
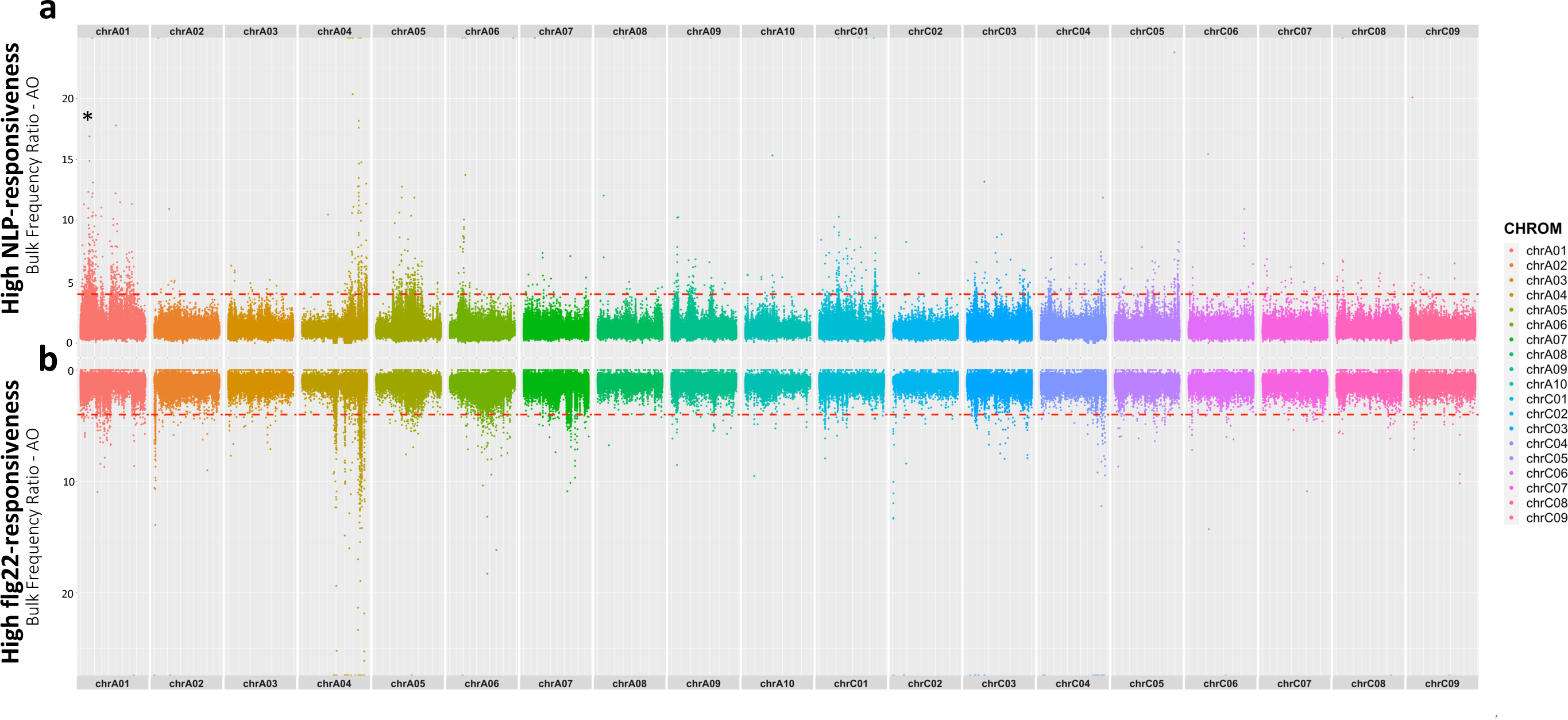
Manhattan plots created with corresponded calculations of the Bulk Frequency Ratios of the variants. **(a)** The highest associated peaks for the High NLP-responsiveness, BFR ratios are calculated between Pool4 and Pool3. 2,676,098 variants (INDELs and SNPs) are plotted along the X-axis on the full genome. **(b)** The highest associated peaks for the High flg22-responsiveness, BFR ratios are calculated between Pool2 and Pool1. INDELs and SNPs are plotted along the X-axis on the full genome (Position of the *BnaA01g02190D* gene on chromosome A01 stated with a star).

**Figure S8.**
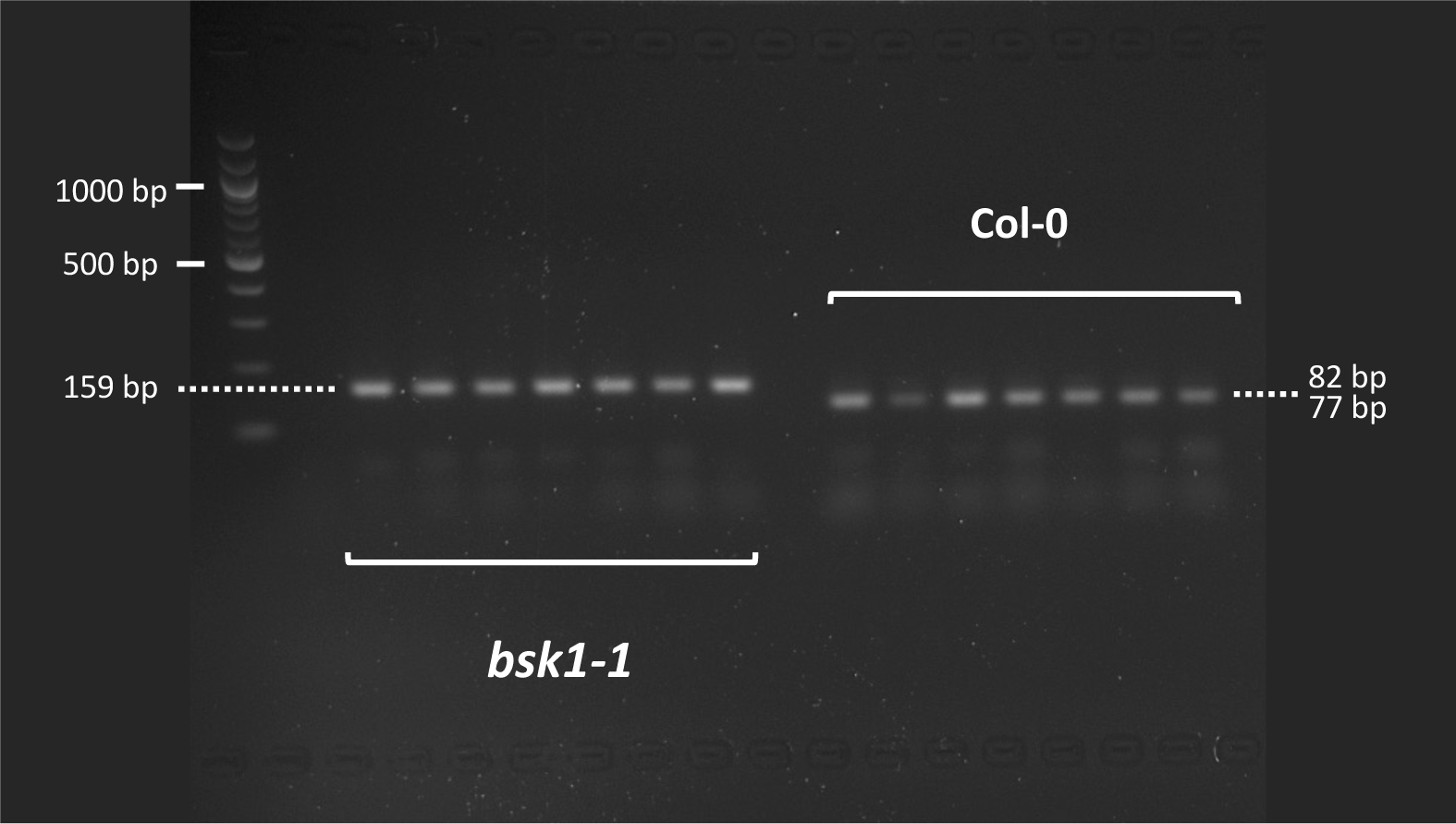
The image of cloned and digested fragments from the mutation region of 8 *bsk1-1* and 8 Col-0 plants. A CAPS marker designed to detect the *bsk1-1* mutation was used to amplify the region of interest. The PCR fragments were obtained and digested with restriction enzyme. As expected, the Col-0 doesn’t have a *bsk1-1* mutation in its genome, so it migrates at 82 and 77 base pair (bp). The *bsk1-1* mutation is confirmed by the fragment, which remains undigested by the enzyme and so migrates at 159bp. The fragments were separated in a 2.4% denatured Agarose gel. 1^st^ row is 100bp DNA Ladder (NEB #B7025). Corresponding sample names of the lanes are represented in the figure.

**Figure S9.**
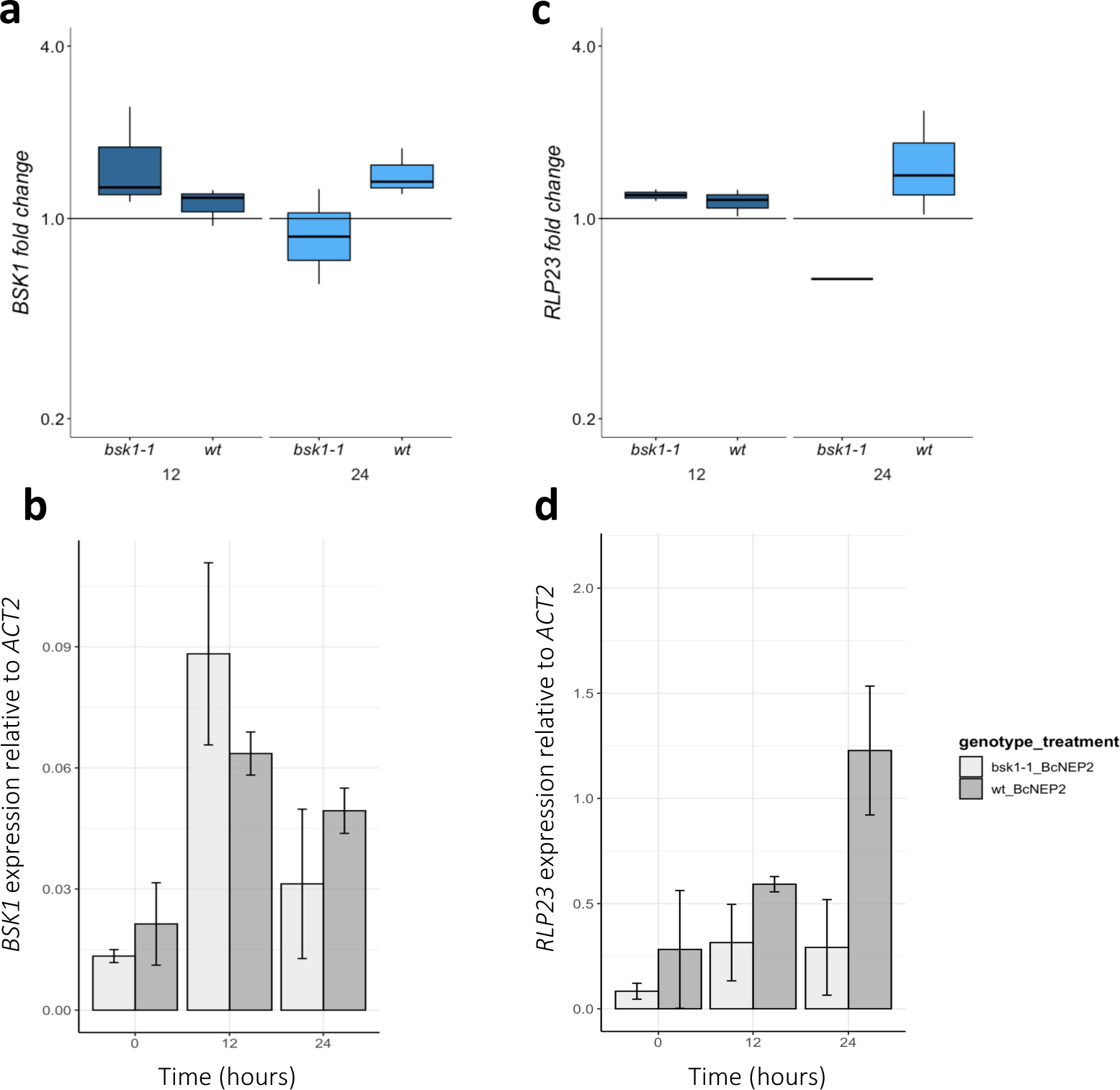
Quantification of induction and expression of *BSK1,* (a,b), *RLP23* (c,d) genes of Arabidopsis after 100nM BcNEP2 treatment at 12 and 24 hours post treatment. mRNA levels of the genes were measured by qRT-PCR in BcNEP2-treated and mock-treated Arabidopsis plants. *ACTIN2* was used as a normaliser. Bars represent means of three biological replicates ± SEM. Asterisks indicate significant differences between expression levels in mock and treated leaves (Student’s t-test *, P <0.05).

